# Deciphering gene regulatory programs underlying functionally divergent naïve T cell subsets

**DOI:** 10.1101/2024.11.06.621737

**Authors:** Hongya Zhu, Ya Jiang, Adrian J. McNairn, Elizabeth A. Fogarty, Cybelle Tabilas, Ravi K. Patel, Jason D. Chobirko, Paul R. Munn, Norah L. Smith, Jennifer K. Grenier, Brian D. Rudd, Andrew Grimson

## Abstract

Naïve CD8+ T cells are a heterogeneous population, with different subsets possessing distinct functions and kinetics upon activation. However, the gene regulatory circuits differentiating these naïve subsets are not well studied. In this work, we analyzed a large collection of public and newly generated RNA-seq and ATAC-seq profiles of different subsets of naïve CD8+ T cells, revealing significant differences in the gene regulatory landscapes between subsets. We leveraged these data by employing a network inference algorithm, Inferelator, to identify the transcriptional regulatory circuits active in each subset. The predicted transcriptional network of the naïve CD8+ T cell pool was validated by multiple orthogonal approaches, including CUT&Tag and Micro-C. Interestingly, our network analysis revealed a novel role for *Eomes* in promoting effector cell differentiation in specific cell subsets. Moreover, we uncovered multiple novel regulators across a variety of subsets and discovered several modules of genes that were co-regulated by shared sets of transcription factors in distinct subsets. Collectively, our data defines the gene regulatory programs differentiating naïve CD8+ T cells and facilitates the identification of novel transcription factors that may alter the propensity of naïve CD8+ T cells to become effector or memory cells after infection.

## Introduction

CD8+ T cells are a core component of the adaptive immune system, protecting the host against intracellular pathogens. Upon stimulation, CD8+ T cells differentiate into short-lived effectors that eliminate infected cells or long-lived memory cells that protect the host against repeat infections^1,2^. In the past, it was thought the naïve CD8+ T cells were homogenous, and cell fates were determined by random factors that arise during priming^3,4^. As a result, a major focus has been on understanding the transcription factors and epigenetic programs that arise after stimulation and drive effector and memory cell differentiation. However, it is now evident that the naïve pool is not uniform but instead comprised of multiple subsets of cells that are pre-programmed to respond in different ways^5–7^. The goal of our study was to identify the gene regulatory programs that underlie the distinct responses mediated by different subpopulations of naïve CD8+ T cells.

A major source of heterogeneity in the antigen-inexperienced pool of CD8+ T cells relates to the amount of homeostatic proliferation that individual cells undergo prior to stimulation^8,9^. The subpopulation of CD8+ T cells that undergo extensive homeostatic proliferation express markers also found on memory cells (CD44, CD122) and are referred to as ‘virtual memory’ (VM) cells^7,10,11^. During infection, VM cells are the first to respond and differentiate into effectors. In contrast, the less proliferative subpopulation of CD8+ T cells (‘true naïve’ or TN cells) express low levels of CD44 and CD122 and respond to infection with slower kinetics.

Another source of heterogeneity in the starting pool of CD8+ T cells corresponds to when individual cells are generated (neonatal life or adulthood)^12,13^. Neonatal CD8+ T cells are generated from fetal liver progenitors and rapidly become terminally differentiated after infection^14,15^. In addition, the neonatal cells are highly responsive to inflammation and can be activated by innate cytokines in the absence of TCR stimulation^16^. Adult CD8+ T cells, in contrast, are produced from adult bone marrow HSCs. Although adult cells are more dependent on TCR signaling than neonatal CD8+ T cells, they have an enhanced ability to from memory and respond to secondary infections. Although previous work has uncovered several transcription Factors (TFs) that are differentially expressed in neonatal versus adult, or TN versus VM cells, such as *T-bet*, *Eomes*, *Runx2* and *Bhlhe40,* the functional implications of these differences are unclear, and the downstream pathways these TFs control have not been elucidated^7,15,17,18^.

In addition to intrinsic factors (homeostatic proliferation, developmental origin), the antigen-inexperienced pool is also shaped by environmental factors, such as microbial exposure. Microbial exposure acting upon thymic progenitors alters the gene expression profiles and functions of CD8+ T cells that are exported to the periphery^19^. Animals that experience high microbial exposure early in development generate ‘dirty’ CD8+ T cells that clear infections more effectively compared to those generated in mice living in a more sterile environment (specific pathogen-free; ‘clean’)^19^. This phenomenon indicates that trained immunity, a concept describing innate immune memory mediated by epigenetic and metabolic reprogramming^20,21^, may be applicable in CD8+ T cells. Indeed, genes characteristic of T cell effector functions are elevated in dirty naïve cells^19^, but how these genes are regulated and what TFs govern such regulatory patterns are unknown.

These different subtypes of naïve CD8+ T cells exhibit distinct kinetics and functions post-activation (Fig. 1A). One grouping, VM, neonatal and dirty cells (fast-acting), respond quickly to infection, while a second comprised of TN, adult and clean cells (slow-acting), respond more slowly. Shared regulatory pathways that program response kinetics within the fast- or slow-acting groups may exist, but this possibility has not been investigated. In contrast, it is possible that gene regulatory programs differ within each of the fast-acting and slow-acting groups, and the identities of such programs are likely to be important in understanding the diversity and complexity of the naïve T cell pool.

**Fig. 1:**
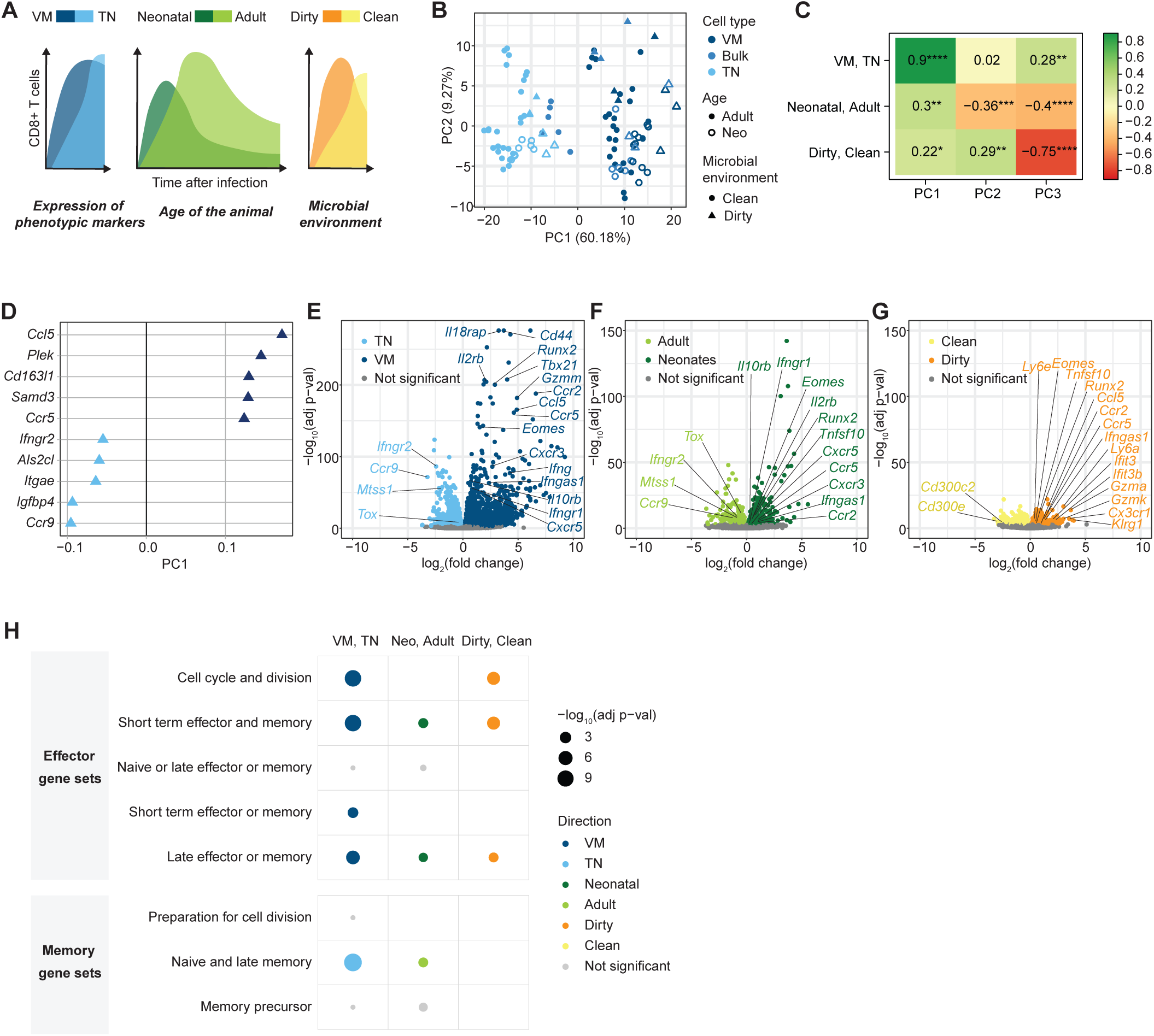
Transcriptional landscapes of naïve CD8+ T cell subsets. (A) Dynamics of naïve CD8+ T cells populations post-activation. (B) PCA of RNA-seq profiles from naïve CD8+ T cell subsets. (C) Pearson correlation, r, between principal components and metadata. ****, P<0.0001; ***, P<0.001; **, P<0.01; *, P<0.05. (D) Top loadings contributing to PC1. (E) Volcano plots showing differentially expressed genes between TN and VM cells, adult and neonatal cells (F), and clean and dirty cells (G). Genes with adjusted p-values larger than 0.001 were plotted as gray. (H) GSEA showing gene sets differentially enriched in different naïve CD8+ T cell subsets.

Transcriptional regulatory network (TRN) inference is a powerful tool that describes the regulatory relationships between transcription factors (TFs) and target genes^22,23^. Constructing cell-type specific TRNs can not only identify key regulators of cell identity and function, but also reveal downstream targets of the regulators, guiding further dissection of these circuits. TRN inference typically relies on expression profiles to link TFs to their targets, assuming that the levels of expression of genes with regulatory relationships are interdependent^23^. State-of-the-art algorithms incorporate chromatin accessibility data to refine predictions, exploiting TF motif occurrence in differentially open chromatin regions to associate TF identifies to target genes^24^. Such approaches combining transcriptional and epigenetic profiling have been shown to be particularly useful in mammalian systems, which possess complex regulatory networks^24,25^.

TRN inference has been employed in understanding key regulators and pathways in many physiological processes^26,27^. To study the transcriptional control of memory CD8+ T cell development, a TRN has been constructed, uncovering novel TFs regulating memory establishment^17^. However, networks for naïve CD8+ T cells have not been characterized. The goal of this study was to generate a TRN in naïve cells, and use this network to understand how variations in gene regulation preprogram divergent phenotypes that manifest post-activation.

To investigate transcriptional regulation in naïve CD8+ T cells, we analyzed a diverse set of RNA-seq and ATAC-seq profiles of naïve CD8+ T cell subsets, and observed substantial differences in the transcriptional and epigenetic landscapes between subsets. We leveraged these data to infer regulatory networks with the network inference algorithm Inferelator, which we selected because of its successful applications in both Innate Lymphoid Cells and T helper 17 cells^24,28^. To validate the naïve CD8+ T cell network, we generated and used a series of orthogonal datasets, including transcription factor binding profiles (CUT&Tag^29^) and genome architecture maps (Micro-C^30^), as well as previously generated transcriptome profiles of T cells with TFs of interest knocked out or over-expressed. Using the network, we identified key TFs in different naïve subsets, which not only revealed previously uncharacterized roles of established regulators, but also implicated novel functions for additional factors such as *Zfp422* and *Thra*. Furthermore, we discovered modules of genes with effector or memory functions that are co-regulated by sets of TFs. Taken together, our work provides a comprehensive understanding of transcriptional programs in naïve CD8+ T cells, revealing novel regulators and regulatory circuits that preprogram naïve cells mediating their divergent kinetics and functions post-activation.

## Results

### Transcriptional landscapes of naïve CD8+ T cell subsets

Naïve CD8+ T cells of different phenotypes, developmental origins or microbial environments exhibit distinct functions and dynamics upon activation^12,14,15,19,31^ (Fig. 1A). The changes in immune responses upon activation are thought to be specified by gene programs established within naïve cells prior to activation, although these programs are not well defined^6,31–34^. To understand the gene expression landscape of naïve CD8+ T cell subsets in a comprehensive manner, we analyzed 102 RNA-seq profiles of naïve CD8+ T cell populations, comprising both public and newly generated data (Fig. 1B, Supplementary Fig. 1A). Principal Component Analysis (PCA) showed that the first component (PC1) separated TN and VM cells, accounting for 60% of the variance across the dataset (Fig. 1B, Supplementary Fig. 1B). Within each TN and VM population, profiles of neonatal and dirty cells were distinct from those of adult and clean cells (Fig. 1B-C, Supplementary Fig. 1C-D). Overall, CD8+ T cells characterized by their ability to respond rapidly to infection (fast-acting cells; VM, neonatal and dirty samples) were located toward the positive end of PC1, implying shared regulatory pathways. Adult versus neonatal cells, and clean versus dirty cells were partitioned further by PC2 and PC3, which accounted for 9% and 6% of the total variance, respectively (Fig. 1C, Supplementary Fig. 1C-G). Top genes contributing to the positive end of PC1 (fast-acting phenotypes) included the *Ccl5*-*Ccr5* chemokine ligand-receptor pair and *Plek* (Fig. 1D)*. Ccl5* is a chemokine important in homing and migration of effector and memory T cells^35^; *Plek* is a target of NFATc1, which controls cytotoxicity^36^. Elevation of the *Ccr5*-*Ccl5* ligand-receptor pair and *Plek* in naïve fast-acting cells is consistent with preprogramming of these cells toward prompt activation and effector fates. The top gene associated with the negative side of PC1 was *Ccr9*, encoding a lineage-associated chemokine receptor selectively expressed in CD8+ T cells^37^ (Fig. 1D). Other genes that drive negative PC1 loadings included *Ifngr2*, which is expressed at lower levels in activated cells^38,39^. Taken together, different naïve CD8+ T cell subsets displayed distinct gene expression patterns, consistent with a preprogramed state underlying divergent post-activation phenotypes.

Differential expression analysis confirmed the striking differences in gene expression between TN and VM cells. Compared to TN cells, VM cells exhibited higher expression of genes encoding phenotypic markers (*Cd44*, *Il2rb*), TFs (*Eomes*, *Tbx21*, *Runx2*), effector molecules (*Gzmm*), and certain chemokines and chemokine receptors (*Ccl5*, *Ccr5*, *Ccr2*, *Cxcr3*, *Cxcr5*, *Il10rb*) (Fig. 1E). Elevated expression of these same genes was also observed in neonatal and dirty cells compared to their respective counterparts, although to a lesser degree (Fig. 1F-G). Gene set enrichment analysis (GSEA) showed that CD8+ T cell effector gene sets were more highly expressed in VM, neonatal and dirty cells compared to their counterparts (Fig. 1H), which likely facilitates their faster response to activation^12,18,19^. Finally, multiple transcription factors showed diverse expression patterns across different naïve CD8+ T cell subsets (Supplementary Fig. 1H, Supplementary Table 1). The distinct expression patterns of many TFs and other genes (Supplementary Fig. 1H-I, Supplementary Tables 1-2) indicate extensive differences in gene regulatory circuits across naïve CD8+ T cell subsets. The substantial changes in the transcriptional landscape of naïve CD8+ T cell subsets underscore the need to identify the TFs that underlie the alternative gene expression programs differentiating naïve CD8+ T cell phenotypes.

### Diverse patterns of chromatin accessibility across naïve CD8+ T cell subsets

To understand the epigenomic landscape of naïve CD8+ T cell subsets, we analyzed 82 newly generated and existing^19,31,34^ profiles of assay for transposase-accessible chromatin with sequencing (ATAC-seq) data^40^, which detects chromatin accessibility, thereby profiling active enhancers, which are enriched at accessible regions. PCA of the chromatin accessibility profiles revealed that PC1, explaining 81% of the variance, separated TN and VM cells (Fig. 2A-B, Supplementary Fig. 2A). Adult versus neonatal, and clean versus dirty cells were split by PC1 to some extent, in combination with PC2 and PC3 (Fig. 2B, Supplementary Fig. 2B-G). The general patterns of chromatin accessibility across the naïve CD8+ cell subsets were similar to those obtained by transcriptome profiling, with PC1 coincident with distinguishing between fast- and slow-responding cells (VM, neonatal, dirty cells versus TN, adult, clean cells) (Fig. 2A-B, 1B-C). Concordant with their expression changes, accessible peaks at promoters of *Ccl5* and *Ccr5* were elevated in fast-acting cells (Fig. 2C-D), while peaks at the promoter and gene body of *Ccr9* were more accessible in slower-acting cells (Fig. 2E).

**Fig. 2:**
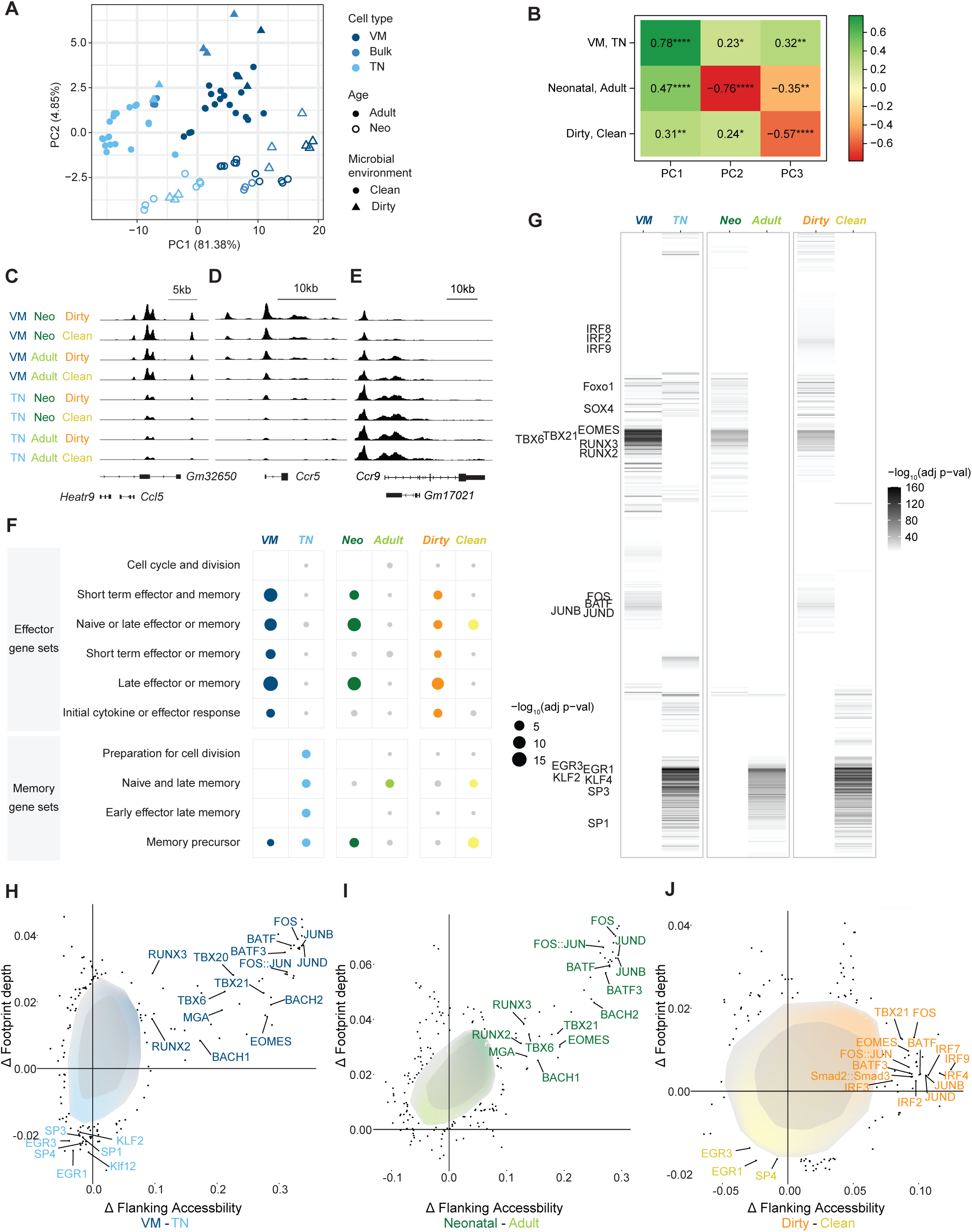
Chromatin accessibility landscapes of naïve CD8+ T cell subsets. (A) PCA of ATAC-seq profiles from naïve CD8+ T cell subsets. (B) Pearson correlation, r, between principal components and metadata. ****, P<0.0001; ***, P<0.001; **, P<0.01; *, P<0.05. (C) Chromatin accessibility at *Ccl5*, *Ccr5* (D) and *Ccr9* (E) loci. (F) Enrichment analysis on genes associated with differentially accessible peaks. (G) TF motif enrichment on differentially accessible peaks. (H) Bag plots showing changes in TF motif footprint depth (y-axis) and flanking accessibility (x-axis) comparing VM and TN cells, neonatal and adult cells (I), and dirty and clean cells (J).

To study chromatin accessibility differences across naïve CD8+ T cell subsets, we identified differentially accessible ATAC-seq peaks between the subset pairs (TN versus VM; adult versus neonatal; clean versus dirty), and associated those regions to the closest genes. Enrichment analysis using gene sets characteristic for CD8+ T cell activation showed that peaks nearby effector genes were more accessible in VM, neonatal and dirty cells compared to their counterparts, while peaks associated with memory genes were more accessible in TN, adult and clean cells (Fig. 2F). This analysis showed that genes with effector and memory functions are regulated epigenetically across naïve subsets. Importantly, these results indicate the existence of gene expression programs shared between fast-acting naïve subsets and also between slow-acting subsets.

Next, we investigated inferred TF activities in naïve CD8+ T cell subsets using the ATAC-seq data. We performed TF motif enrichment on differentially accessible peaks of naïve CD8+ T cell subsets, assuming that a TF with high motif occurrences on subset-specific enhancers would have higher activity in that subset. We observed overlapping motif enrichment patterns within the fast- or slow-responding groupings (Fig. 2G). Peaks more accessible in fast-acting cells had enriched motifs for TFs in the TBX and RUNX families, including Eomes, T-bet and Runx2, which regulate CD8+ T cell effector and memory functions. Motifs of AP-1 factors (Fos-Jun) and BATF were enriched in VM and dirty cells compared to their counterparts, which control adaptive effector programs as well as bystander activity^16,41,42^. Moreover, motifs corresponding to the IRF family were more enriched in dirty cells compared to clean; TFs in this family play important roles in CD8+ T cell effector differentiation, proliferation and cytotoxic functions^43,44^. In contrast, TFs implicated as being more active in the slow-acting group include EGR, KLF and SP family members. KLF2 and KLF4 have been suggested to maintain naïve T cell quiescence^45^, and Egr2 and Egr3 operate as a checkpoint controlling clonal expansion and effector differentiation^46^. This analysis demonstrated that motifs of TFs promoting effector functions were enriched at accessible enhancers enriched in the fast-responding subsets, whereas motifs of TFs repressing effector programs were enriched in those of the slow-acting subsets.

To assess TF activity in a more rigorous manner, we conducted BaGFoot (Bivariate Genomic Footprinting) analysis, which examines motif flanking accessibility while also quantifying TF footprint depth as a proxy for binding^47^. This analysis revealed that TFs in TBX, RUNX, AP-1 and BATF families (including Eomes, T-bet, Runx2, Fos-Jun, Batf) had more and deeper footprints in VM, neonatal, and dirty cells compared to their counterpart subsets, indicating higher activities of those TFs in the fast-acting cells (Fig. 2H-J). Enhanced binding of AP-1 factors contributes to the increased innateness of neonatal CD8+ T cells^16^. Moreover, TFs in IRF families had higher activity in dirty cells compared to clean (Fig. 2J); TFs in EGR and SP families were more active in TN and clean cells (Fig. 2H, J). These results recapitulated and extended those of TF motif enrichment, suggesting that the identified TFs were not only associated with accessible enhancers but also bind the same sets of enhancers. Taken together, these analyses indicate substantial epigenetic differences between naïve CD8+ T cell subsets.

### A transcriptional regulatory network for naïve CD8+ T cells

To identify transcriptional regulatory circuits active in different naïve CD8+ T cell subsets, we used Inferelator, an established and powerful transcriptional regulatory network inference algorithm designed to leverage RNA-seq and ATAC-seq data^24^. The algorithm uses expression profiles across diverse cell types to model gene expression patterns and predict TF-gene interactions, while using chromatin accessibility coupled with TF motif analysis to refine the predictions^24^. Using RNA-seq and ATAC-seq profiles of naïve CD8+ T cells as input to Inferelator, we constructed a transcriptional regulatory network for naïve CD8+ T cell subsets, comprising 602 TFs and 8,186 genes (Fig. 3A). Among the 65,616 predicted TF-gene regulatory interactions, 83% were supported by TF motif occurrence in proximal chromatin accessible regions. The predictions were robust against different network construction parameters (Supplementary Fig. 3A-C).

**Fig. 3:**
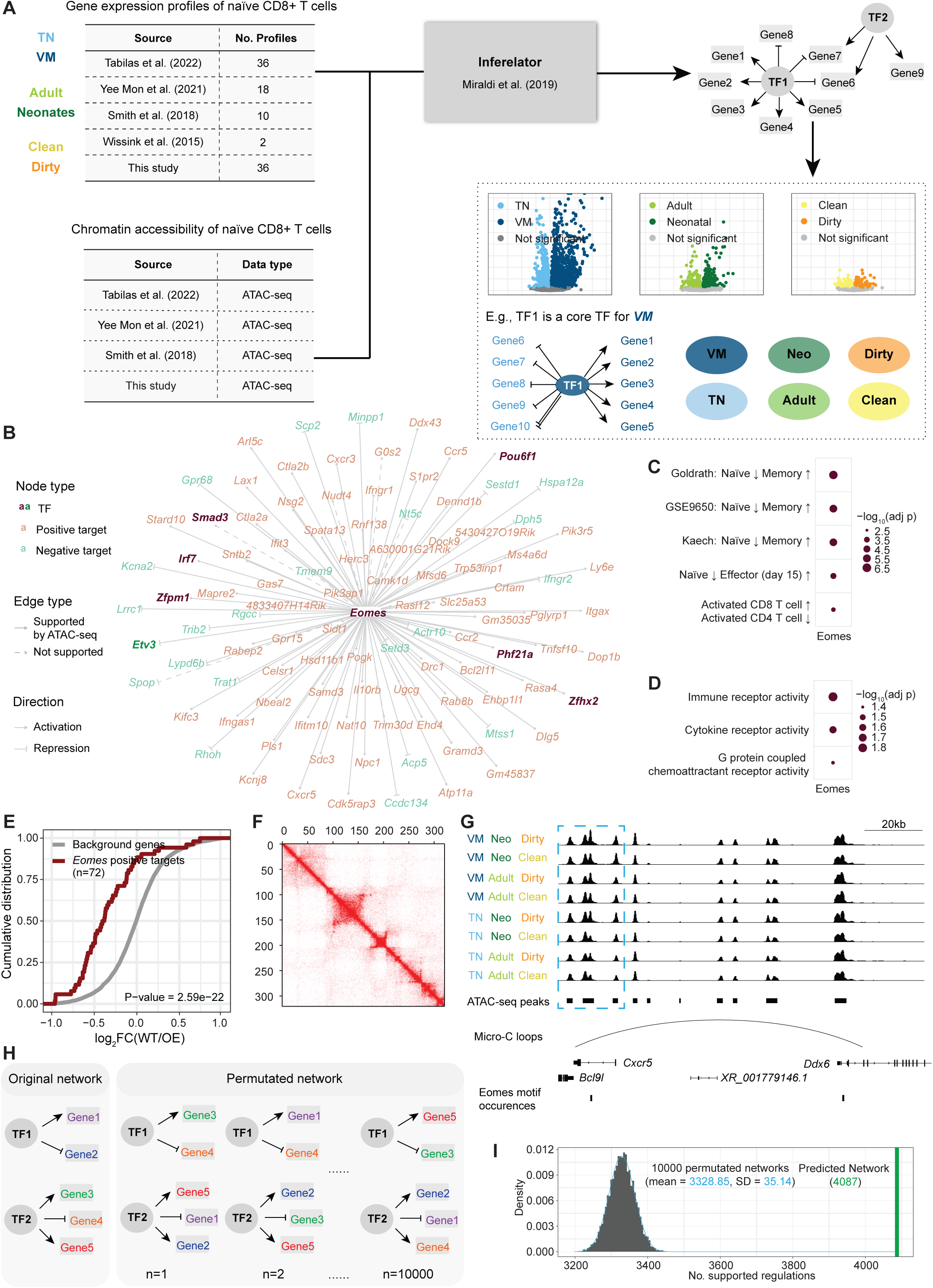
A transcriptional regulatory network for naïve CD8+ T cells. (A) Schema describing construction of transcriptional regulatory network in naïve CD8+ T cell populations. (B) Predicted targets of *Eomes* in the network. (C) Hypergeometric test on *Eomes* positive targets using ImmuneSigDB gene sets, showing gene sets describing states and perturbations of CD8+ T cells. (D) Hypergeometric test on Eomes positive targets using GO Molecular Functions gene sets. (E) Cumulative distribution plot showing significant differences in distributions of log_2_FC (WT/*Eomes* OE; x-axis) between predicted positive targets of *Eomes* and background genes (Wilcoxon test). (F) Example of Micro-C data at location 4M-12M on Chromosome 4 with resolution 25K. (G) Chromatin accessibility of *Cxcr5* and a distal enhancer connected by a Micro-C loop, with *Eomes* motif occurrences. (H) Schema depicting shuffling (10,000 shuffles) predicted network while maintaining network properties. (I) Number of TF-gene regulatory interactions shared by the Micro-C dataset and the original or permutated networks.

We exemplify the predicted network with a subnetwork centered around *Eomes* (Fig. 3B), a master regulator for effector and memory functions in CD8+ T cells^48^. *Eomes* is expressed at higher levels in all fast-acting cells (VM, neonatal and dirty) compared to their counterparts (Fig. 1E-G). Among the 102 predicted *Eomes* targets, 92 had *Eomes* motifs at nearby enhancers. Among the 79 target genes predicted to be activated by *Eomes*, 37 had significantly higher expression levels in at least two fast-responding cell types, including genes encoding cytokines, chemokine receptors and immune regulators (*Tnfsf10*, *Ccr5*, *Ccr2*, *Cxcr3*, *Cxcr5*, *Ifngr1*, *Il10rb*, *Ifngas1*, *Ly6e*) (Fig. 1E-G). Targets of *Eomes* have been identified in exhausted CD8+ T cells^49^ and natural killer cells^50^ by integrating binding profiles and RNA-seq following *Eomes* overexpression or inhibition. Comparing our predicted *Eomes* targets to those identified elsewhere, we observed multiple overlaps (e.g., *Ccr5*, *Ctla2b*, *Cxcr3* and *Samd3*), supporting our predictions. This overlap is notable considering that the validation data is derived from non-CD8+ T cells, and cell type specificity in TF binding and regulation is common. Indeed, our predicted *Eomes* targets also include many genes not identified in these other cell types (e.g., *Cxcr5*, *Ly6e*, *Ifngas1*). GSEA on predicted positive targets of *Eomes* showed enrichment of memory and effector genes, concordant with the established role of *Eomes* (Fig. 3C). *Eomes* predicted targets were also enriched in cytokine receptors (Fig. 3D), suggesting that Eomes may also function by modulating certain naïve subsets’ ability to receive cytokine signals. Comparable subnetworks exist for additional TFs such as *Foxo1*, *T-bet* and *Bach2* (Supplementary Fig. 3D-F).

To evaluate the *Eomes* predictions systematically, we overlayed the predicted targets on an *Eomes*-overexpression (OE) RNA-seq dataset from CD8+ single positive cells^51^, data not employed in the network inference process. The predicted positive targets of *Eomes* were expressed at higher levels when *Eomes* was overexpressed, compared to background genes (Fig. 3E). In contrast, the predicted negative targets of *Eomes* had lower expression levels (Supplementary Fig. 3G). We observed more predicted positive targets of *Eomes* than negative (79 versus 23), consistent with findings suggesting that Eomes is typically an activating TF but can function as a repressive factor^49,50^. Similar results were obtained when we evaluated predicted regulations of *Foxo1*, a TF essential for CD8+ T cells memory function^52^. Using RNA-seq profiles of naïve CD8+ T cells from a *Foxo1* knockout model^53^, we observed that *Foxo1* predicted positive targets were lower in expression in *Foxo1* knockout samples compared to background, with negative targets showing the opposite trend (Supplementary Fig. 3H-I). These analyses demonstrated that the expression levels of predicted *Foxo1* and *Eomes* targets validate using independent and orthogonal data, suggesting that the overall network successfully captures *in vivo* gene regulatory networks.

To evaluate the network more broadly, we performed Micro-C^30^, a derivative of Hi-C technology designed to capture genome structure at fine scale, on naïve CD8+ T cells (Fig. 3F). Overlapping Micro-C identified loops (e.g., enhancer-gene interactions) with ATAC-seq peaks, we identified open chromatin loops in naïve CD8+ T cells, and derived a set of potential TF-gene regulatory interactions using the loops. Specifically, we connected TFs with motif occurrences on one end of the loop to genes close to either loop end. As an example of distal enhancers identified by Micro-C, we examined an *Eomes* predicted target, *Cxcr5*. With increased accessibility observed at its promoter in the fast-responding cells (VM, neonatal and dirty), we found a putative enhancer that contains an *Eomes* motif linked by a Micro-C loop to the *Cxcr5* promoter (Fig. 3G). To calculate the significance of overlap between our network and Micro-C derived TF-gene interactions, we performed a permutation test by shuffling the network. We generated 10,000 random networks by reordering the target genes, so that network properties (numbers of nodes and edges, number of targets regulated per TF and number of TFs regulating each gene) were maintained (Fig. 3H). Overlaps between the random networks and the Micro-C derived linkages were then determined and compared to the predicted network. One caveat of this analysis is that our Micro-C experiment was conducted using a pool of naïve cells from clean adults. Therefore, regulatory interactions predicted for neonatal and dirty cells are unlikely to validate using this approach. In addition, the Micro-C derived TF-gene regulations were generated based on TF motif occurrences, which are expected to be noisy with many false positives^54^). Nonetheless, we observed significant overlap (p < 10^-5^) between the Micro-C dataset and our network, compared to overlaps with the random networks (Fig. 3I). Thus, our network, which is predicted to capture consequential TF-gene interactions, is significantly enriched in physical TF-gene linkages identified by Micro-C.

### Core TFs in different naïve CD8+ T cell subsets

We next sought to identify specific TFs that our network implicated as key factors responsible for the divergent properties of different subsets. We took a two-step approach, inspired by previous studies^24^: first, we used differential expression analysis to identify genes more highly expressed in each subset (Fig. 1E-G); second, we performed GSEA on predicted targets of each TF using the differential expression results. As an example, TFs that specify VM cell specific gene regulation are expected to activate genes more highly expressed in VM subsets, or repress genes more highly expressed in the alternative subset: TN. Such a pattern would identify TFs that promote expression of genes elevated in VM cells, and/or repress genes in VM cells that are expressed highly in TN cells; these TFs may contribute to establishment or maintenance of the VM identity – we refer to them as ‘core’ TFs for VM cells. Using this approach, we identified core TFs for each of the naïve CD8+ T cell subsets.

Among core TFs identified for VM identity (Fig. 4A), we found TFs with established roles in T cell development and function, including *c-Myb*^55^, *Bhlhe40*^56^, *Bach2*^41^ and *Tcf3*^57^. TFs with roles in CD4/CD8 lineage choices were also prominent, such as *Runx1*, *Runx3*^58^ and *Patz1*^59^. *Eomes*, critical for VM cell formation^11^, was recovered as a core TF for VM cells. TFs with roles in proliferation and cell cycle progression (*Foxm1*, *E2f1*, *E2f2*, *E2f7* and *E2f8*) were also identified as VM core TFs (Fig. 4B); VMs are more proliferative than TN cells after antigen stimulation^7,10^. In contrast, TFs known to restrict CD8+ T cell differentiation or promote naivety and quiescence were identified as core TFs for TN cells; for example, *Ikzf1*^60^, *Lef1*^61^ and *Foxo1*^52^. Overall, the majority of the core TFs we defined are supported by the literature. In particular, *Eomes*, an established master regulator of CD8+ T cell differentiation and function^48,62,63^, was predicted through our network analysis to promote effector functions in neonatal cells.

**Fig. 4:**
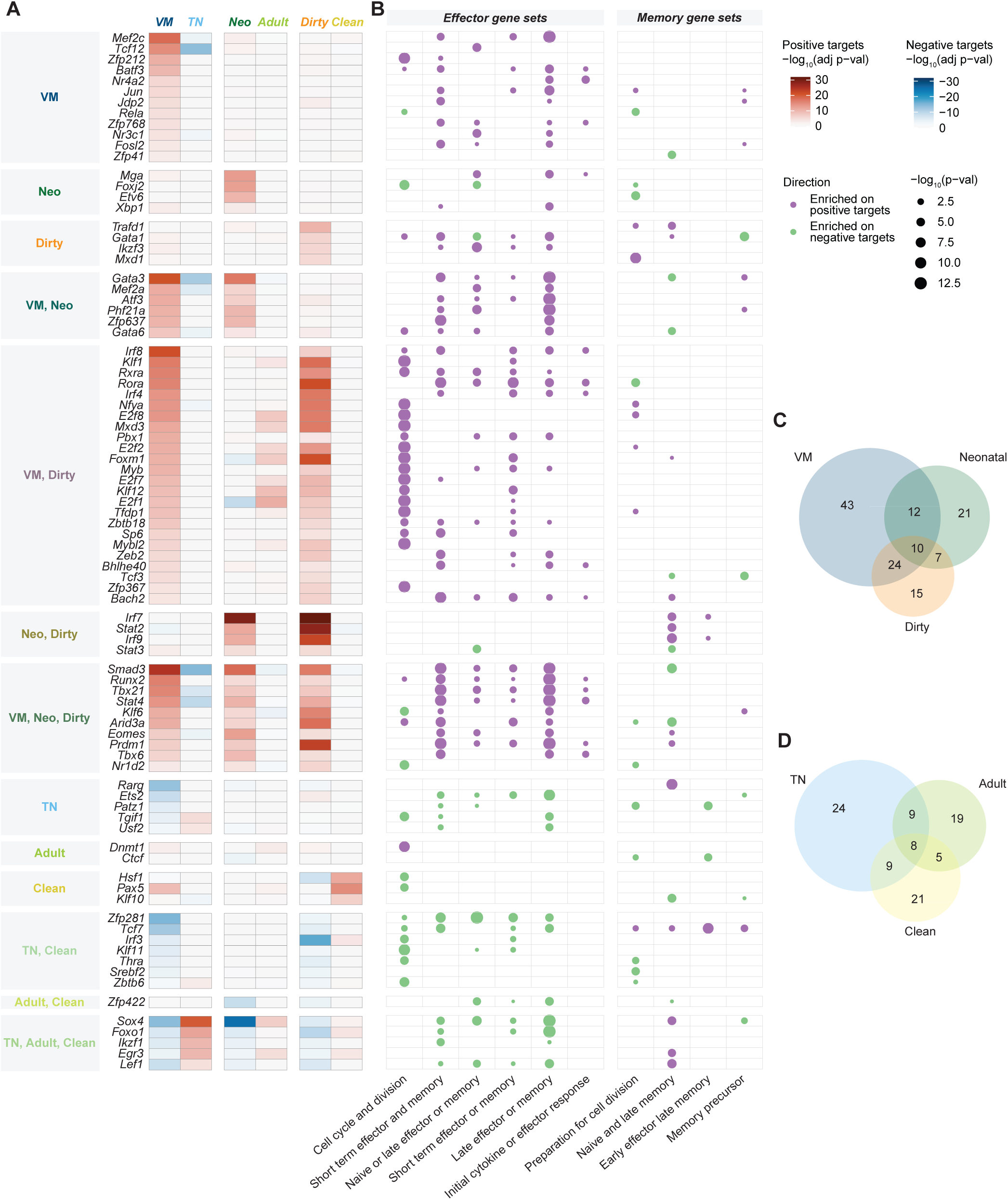
Core TFs in different naïve CD8+ T cell subsets. (A) GSEA of predicted targets of core TFs using differential expression of genes between different pairs of naïve CD8+ T cell subsets, showing core TFs with at least 1 ImmGen gene set enrichment. (B) Hypergeometric test on predicted targets of core TFs using gene sets from ImmGen delineating CD8+ T cell immune response. P-values smaller than 10^−13^ were plotted as 10^−13^. (C) Core TFs shared by VM, Neonatal and Dirty cells. (D) Core TFs shared by TN, Adult and Clean cells.

Importantly, we discovered multiple core TFs shared across the fast-responding cells (VM, neonatal and dirty), suggesting overlapping regulatory circuits between them (Fig. 4A-C, Supplementary Fig. 4A). Several of the fast-responding core TFs have established roles in CD8+ T cell effector functions, including *Eomes*, *Tbx21* and *Prdm1*. The role of *Tbx21* is well-established in control of effector CD8+ T cell generation^64^, and *Prdm1* is known to promote terminal differentiation in effector CD8+ T cells^65^. Furthermore, *Irf8*, a core TF for VM and dirty subsets, has validated roles in driving CD8+ T cell effector differentiation^43^. GSEA using gene sets typifying CD8+ T cell response suggested that these TFs activate effector genes, recapitulating their roles in promoting effector functions (Fig. 4B, Supplementary Fig. 4B). This analysis demonstrated that TFs predicted to establish or maintain VM, neonatal and dirty cell types can drive effector programs, likely preparing them for prompt activation upon infection^11,14,19^.

Overlap between the TN, adult and clean core TFs was also observed (Fig. 4A, D, Supplementary Fig. 4C). *Foxo1*, *Egr3* and *Sox4* were identified as core TFs shared by the slow-acting subsets, all with established roles in opposing effector programs or regulating memory development^17,46,52^. Confirming this pattern, GSEA showed that TFs important for the slow-responding subsets were predicted to repress effector genes and activate genes associated with memory functions (Fig. 4B, Supplementary Fig. 4D). This analysis showed that TFs repressing effector programs play important roles in TN, adult and clean cells, potentially contributing to the slower immune response characteristic of these cells^11,14,19^.

Our approach also uncovered novel TFs without established roles in CD8+ T cells (Fig. 4A, B). Among the TFs identified to be important in the fast-responding subsets, *Mef2a*, a TF with roles in myocardial development^66,67^, was identified as a VM and neonatal core TF, predicted to activate effector genes. *Tfdp1*, encoding a protein that heterodimerizes with E2F proteins and regulates the cell cycle^68^, was predicted to promote cell cycle gene sets in VM and dirty cells, which are highly proliferative. Furthermore, several novel TFs were identified as core TFs for the slow-responding subsets. *Thra*, encoding a hormone receptor, was identified as a TN and clean core TF, which we predicted to repress cell cycle genes (Fig. 4B). *Zbtb6* was also identified as a TN and clean core TF, with many members in the same family known to play roles in T cell development and functions^69^. Finally, *Zfp422*, encoding a zinc finger protein with roles in skeletal muscle differentiation^70^, was identified as a core TF for adult and clean cells, with predicted functions in repressing effector genes. Identification of these novel TFs can help better understand the establishment of fast- and slow-responding cells and the regulation of CD8+ T cell activation in naive cells.

### *Eomes* binding at effector genes in neonatal cells

To validate the predicted functions of *Eomes* in promoting effector functions in neonatal cells, we generated its binding profiles by performing *Eomes* CUT&Tag^29^ in naïve CD8+ T cells from adult wildtype and adult Lin28b transgenic mice, which reprogram hematopoietic stem cells to phenocopy the fetal progenitors, generating CD8+ T cells that recapitulate all phenotypes characteristic of neonatal cells ^15,71–74^. Motif enrichment analysis on *Eomes* CUT&Tag peaks revealed enrichment of T-box family motifs (Fig. 5A). We identified the CUT&Tag-derived *Eomes* targets through associating peaks to nearest genes. The genes proximal to *Eomes* binding sites exhibited significant enrichment in gene sets related to CD8+ T cell effector and memory functions (Supplementary Fig. 5A), congruent with *Eomes*’s role as a master regulator of CD8+ T cell immune response. Pronounced enrichment of gene sets pertaining to the immune system and cytokine signaling were also observed, reinforcing the hypothesis that *Eomes* modulates expression of cytokine receptors (Fig. 5B, 3D). Importantly, we detected significant overlap between *Eomes* targets derived from CUT&Tag and those predicted through our network analysis (Fig. 5C), with additional overlap when considering only CUT&Tag peaks containing the *Eomes* motif (Supplementary Fig. 5B).

**Fig. 5:**
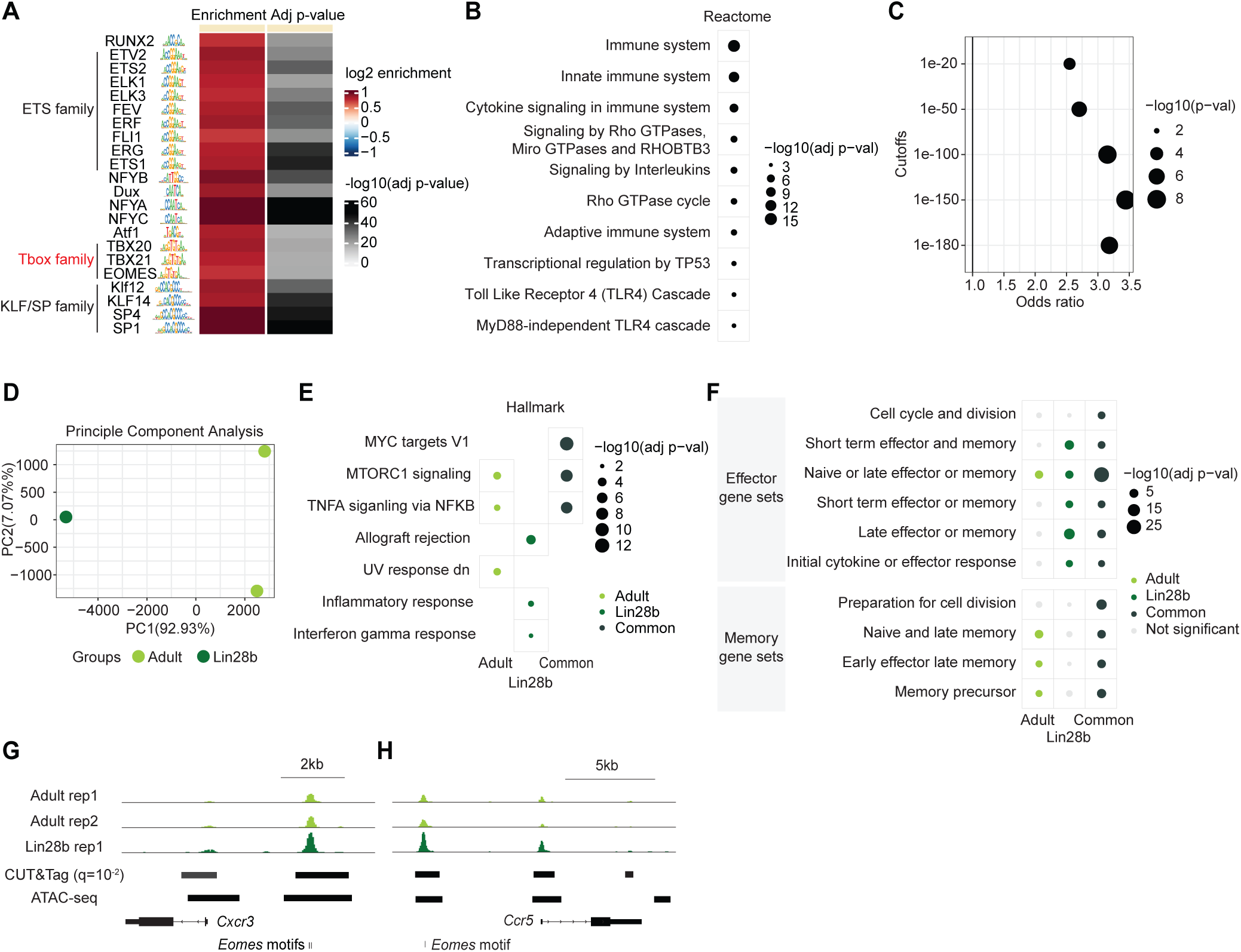
*Eomes* binding at effector genes in neonatal cells. (A) TF motif enrichment on *Eomes* CUT&Tag peaks, showing motifs with adjusted p-values < 10^−15^ and log_2_ enrichment > 0.7. Adjusted P-values smaller than 10^-50^ were plotted as 10^-50^. (B) Enrichment analysis on genes associated with *Eomes* CUT&Tag peaks using Reactome gene sets. (C) Enrichment of CUT&Tag-derived *Eomes* targets and network-derived *Eomes* targets at CUT&Tag peaks of different cutoffs. (D) PCA of CUT&Tag profiles from naïve CD8+ T cells from adult and Lin28b mice. (E) Enrichment analysis on genes associated with peaks with enhanced binding in adult cells, Lin28b cells or common to both cell types, respectively, using Hallmark and ImmGen gene sets (F). (G) CUT&Tag peaks in adult and Lin28b cells of *Cxcr3* and *Ccr5* (H).

PCA revealed notable differences between the *Eomes* binding profiles in naïve adult and Lin28b (neonatal-like) CD8+ T cells (Fig. 5D), with peaks specific to adult (15% of the total) and Lin28b (5%) cells identified (Supplementary Fig. 5C). To examine differential binding patterns of *Eomes*, we mapped the adult- or Lin28b-specific peaks to their nearest genes, and conducted enrichment analysis to explore their functional roles. We found that genes proximal to Lin28b-specific peaks were implicated in effector responses, while those near adult-specific peaks were involved in memory functions (Fig. 5E-F). Genes located near shared peaks were enriched for both effector and memory functions (Fig. 5E-F). *Cxcr5* and *Ccr5* were identified as *Eomes* targets in our predicted network (Fig. 3B). *Eomes* binding and motif occurrences were detected at peaks proximal to both genes, with enhanced binding in Lin28b cells compared to adult cells (Fig. 5G-H). Thus, *Eomes*, a master regulator of CD8+ T cell immune responses, binds to both effector and memory genes in naïve CD8+ T cells. The enhanced binding of *Eomes* to effector genes in Lin28b cells supports our prediction that *Eomes* drives effector functions in neonatal naïve cells.

### Co-regulatory TF modules preprogram naïve cells for effector and memory functions

Genes with coordinated functions are often co-regulated by common TFs^75,76^. To identify TF sets s that function together, we grouped TFs into co-regulatory modules based on overlap of their positive targets^24^. Several modules were predicted to promote effector functions, based on GSEA of their shared targets (Fig. 6A-B, Supplementary Fig. 6A). Among them, one module, comprised of *Stat4*, *Runx2* and *Zfp367*, was predicted to promote the fast-acting phenotypes (VM, neonatal and dirty) and effector functions (Fig. 6C). *Stat4* is required for innate cytokine responsiveness of CD8+ T cells^77^ and cytotoxicity of CD8+ T cell in islets^78^. *Runx2* possesses roles in formation of long-term memory in CD8+ T cells^79^. *Zfp637*, with roles in inhibiting muscle differentiation^80^ and preventing cell senescence^81^, has not been associated with T cell functions. Our analysis suggested that *Zfp637* may function with *Stat4* and *Runx2*, promoting both effector (Fig. 6A-B) and memory functions (Supplementary Fig. 6B), with multiple genes predicted as positive targets of at least two TFs (Fig. 6C), including two serine proteases (*Gzmb*, *Gzmm*) and *F2r*, a thrombin receptor required for CD8+ T cell cytotoxicity^82^. Binding motifs for all three TFs occurred close by *F2r*, with a gradual increase of chromatin accessibility from slow-(TN, adult, clean) to fast-responding (VM, neonatal, dirty) cells (Fig. 6D). The accessibility changes indicated stronger activation of *F2r* by these three TFs in the fast-responding cells. This novel TF module, composed of *Stat4*, *Runx2* and *Zfp367*, may promote fast-responding phenotypes and effector functions through driving expression of *F2r* and other genes.

**Fig. 6:**
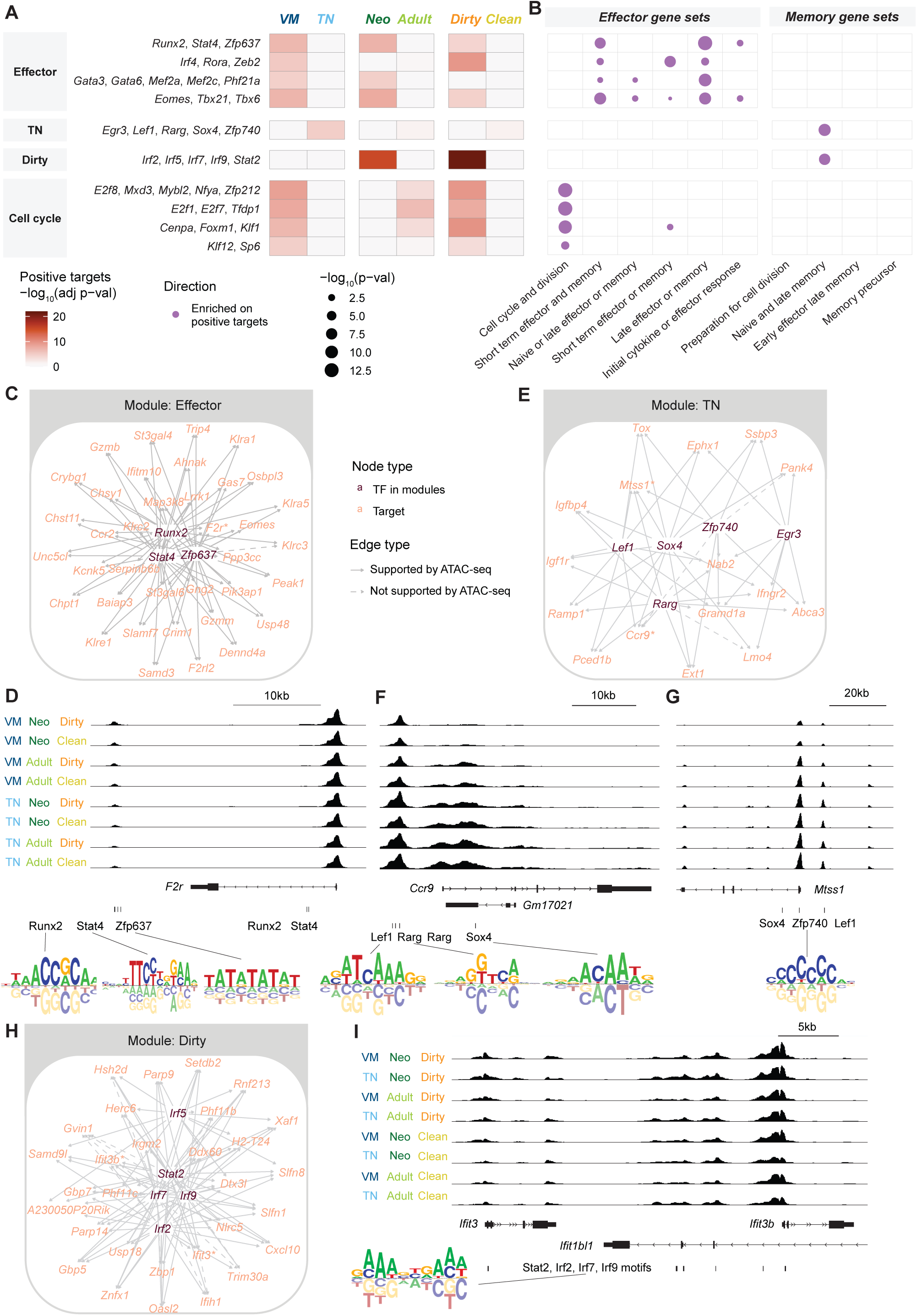
Co-regulatory TF modules preprogram naïve cells for effector and memory functions. (A) GSEA of predicted targets shared by >50% of TFs in each TF module, showing modules with at least 1 ImmGen gene set enrichment. (B) Hypergeometric test on predicted targets of core TFs using ImmGen gene sets delineate CD8+ T cell immune response. P-values smaller than 10^-13^ were plotted as 10^-13^. (C) Subnetwork of *Runx2-Stat4-Zfp637* module, showing TFs and targets shared by at least two TFs. (D) Chromatin accessibility of *F2r* and example TF motifs. (E) Subnetwork of *Egr3*-*Lef1*-*Rarg*-*Sox4*-*Zfp740* module, showing TFs and targets shared by more than 3 TFs. (F) Chromatin accessibility of *Ccr9* and TF motifs. (G) Chromatin accessibility at *Mtss1* locus and motifs of predicted TFs in the module. (H) Subnetwork of *Irf2*-*Irf5*-*Irf7*-*Irf9*-*Stat2* module, showing TFs and targets shared by more than 4 TFs. (I) Chromatin accessibility around *Ifit3* and *Ifit3b* loci and motifs of predicted TFs in the module.

An additional module of interest was comprised of core TFs for TN cells (*Egr3*, *Lef1*, *Rarg*, *Sox4* and *Zfp740*) (Fig. 6E). *Egr3*, *Lef1* and *Sox4* inhibit CD8+ T cell effector differentiation^46^, promote memory development^17^, and maintain T cell identity^83^, respectively. The role of *Zfp740*, on the other hand, has been implicated but not well studied in CD8+ T cells^84^. GSEA on shared targets of this module showed enrichment of naïve and late memory gene sets (Fig. 6B). *Ccr9*, the top gene driving PCA loadings of the TN cells (Fig. 1D), was a shared target of this TF module with motif occurrences at its promoter region (Fig. 6F). *Mtss1* (Metastasis suppressor protein 1), with roles in repressing cancer progression^85^, was also identified as a shared target of this module (Fig. 6E). We observed higher expression of *Mtss1* in TN cells (Fig. 1E) and increased accessibility at its promoter (Fig. 6G), with binding motifs corresponding to *Lef1*, *Sox4* and *Zfp740* at the locus. These analyses suggest that this TF module promotes TN identity and helps maintain effector genes in a repressed state.

We also identified a module containing core TFs for dirty cells (*Irf2*, *Irf5*, *Irf7*, *Irf8* and *Stat2*; Fig. 6A, H). GSEA of their shared targets showed enrichment on gene sets containing memory genes, and interferon alpha and gamma response genes (Fig. 6B, Supplementary Fig. 6B-C). *Ifit3* and *Ifit3b*, encoding Interferon-induced antiviral proteins, were among the shared targets of the TF module, and these interactions are supported by increased chromatin accessibility in dirty cells at promoters of these genes (Fig. 6H-I). Finally, four modules promoting cell cycle and division were uncovered (Fig. 6B, Supplementary Fig. 6D). TFs in those modules also emerged as core TFs for VM cells (Fig. 6A), consistent with the increased proliferative capacity of VM cells^7^. GSEA showed enrichment on G2M checkpoint, mitotic spindle, microtubule motor activity and effector genes (Supplementary Fig. 6B-C). Our analyses of these co-regulated modules can advance our understanding of how multiple TFs acting in concert govern regulatory pathways in different naïve CD8+ T cell subsets.

## Discussion

We investigated transcriptional regulation in naïve CD8+ T cell subsets, detailing regulation that establishes cell states with differential activation kinetics and effector/memory potential. We focused on naïve cells of different phenotypes (VM and TN), developmental origins (adult and neonatal), and under different microbial environments (clean and dirty). As the first study to examine a wide range of naïve subsets, our targeted yet comprehensive scope provided substantial analytical power. We revealed extensive differences in the transcriptional landscapes of naïve CD8+ T cell subsets, with elevated expression of effector genes in VM, neonatal and dirty cells compared to their counterparts, which likely contributes to their enhanced effector functions upon stimulation. Furthermore, we observed substantial epigenetic differences across naïve CD8+ T cell subsets, including changes in chromatin accessibility of effector and memory genes and activities of TFs regulating T cell naivety, memory potential and differentiation. These differences indicated the role of epigenetic priming in establishing the distinct post-activation kinetics between naïve CD8+ T cell subsets.

Leveraging RNA-seq and ATAC-seq profiles, we constructed a transcriptional regulatory network for naïve CD8+ T cells using a powerful network inference algorithm, Inferelator^24^. We validated the network using TF perturbation RNA-seq profiles and a Micro-C dataset in naïve CD8+ T cells. Through analyzing predicted TF targets, we identified core TFs in each naïve subset. We revealed novel functions for established regulators, such as the role of *Eomes* in promoting effector functions in neonatal cells. Eomes exhibited enhanced binding proximal to effector genes in neonatal-like cells, corroborating our predictions. Novel TFs without previously described roles in CD8+ T cells were also implicated, including *Mef2a*, *Tfdp1*, *Thra*, *Zbtb6* and *Zfp422*. Enrichment analysis on their targets suggested roles in control of effector programs. By examining shared targets, we identified modules of TFs likely functioning in the same pathway. For example, we uncovered one TF module (*Egr3*, *Lef1*, *Rarg*, *Sox4* and *Zfp740*) with potential roles in repressing effector functions, pronounced in TN cells.

Our study provided a comprehensive map of transcriptional regulation for several non-conventional naïve subsets: those bearing activation markers while being antigen inexperienced (VM cells), produced in early life (neonatal cells), and generated in animals under early microbial exposure (dirty cells). While prior studies have examined aspects of regulation in these cells^19,31,33,51,86,87^, we offered a more extensive and in-depth analysis with broader scope, including novel insights into the TFs and regulatory commonalities between different naïve subsets. Our unbiased approach identified *Eomes* as a key TF promoting and establishing VM and neonatal cells, as previously described^7,31,51^, and more importantly, uncovered multiple novel TFs with enhanced activities in those cell types, indicating their previously unrecognized roles. We highlighted multiple TFs driving the phenotype of dirty cells, an underexplored area, thus bridging a significant gap in the field.

Both VM and neonatal CD8+ T cells mediate bystander protective immunity by responding to cytokines in a TCR-independent manner^16,86,88^. We observed elevated expression of chemokine receptors in VM and neonatal cells, aligned with their increased cytokine responsivity. The gene regulation underlying bystander activation of neonatal CD8+ T cells was previously associated with BACH2 and AP-1 TFs^16^. Increased footprints of those TFs were observed in both VM and neonatal cells, suggesting a shared mechanism enabling bystander activity. Dirty cells generated from animals experiencing early microbial exposure also exhibited enhanced binding of such TFs and elevated expression of cytokine receptors, indicating that they may also be capable of this enhanced innate-like function. Future research investigating bystander activation in cells under the impact of microbial environments will be required to test this hypothesis.

Our analyses uncovered shared regulatory patterns within the fast-acting naïve subsets (VM, neonatal and dirty) as well as within the slow-acting ones (TN, adult and clean). Concordant changes in gene expression signatures, chromatin accessibility patterns and TF activities were observed, indicating common regulatory circuits in the fast- and slow-acting groups. It is reasonable to infer that the shared regulatory pathways pave the way for their common functional traits. VM, neonatal and dirty cells are generated from different progenitors or under different developmental conditions: VM cells are generated in response to homeostatic processes^7^; neonatal cells are derived from fetal progenitors^12^; dirty cells are generated by animals under early microbial exposure^19^. These results suggest questions for further exploration, for example, elucidating the environmental cues or signaling pathways activating the shared core TFs and regulatory circuits in these naïve subsets.

Our data identified multiple novel TFs without implicated roles in T cell functions. *Mef2a* is known to play a crucial role in myocardial development^66,67^, and promotes proliferation in coronary artery endothelial cells^89^ and tumor progression in colorectal cancer^90^. In the innate immune system, *Mef2a* regulates inflammatory responses in macrophages^91^. Our analyses suggest a novel role of *Mef2a* in T cell functions – to promote fast-responding subsets and activate effector genes, possibly through activating inflammation-related genes^91^. *Tfdp1* regulates the cell cycle and E2F proteins^92^, and has been recently implicated in modulating global chromatin accessibility^93^. Our analysis suggests the novel role of *Tfdp1* in promoting fast-responding subsets through activating cell cycle genes, potentially contributing to the specification of the gene expression program of VM and dirty cells.

Chemokine signaling is fundamental for directing immune responses^94^. We observed diverse expression patterns of cytokine receptors in different naïve CD8+ T cell subsets. *Ccr2*, capable of regulating cytokine expression profiles and T cell inflammatory response^95^, was more highly expressed in VM, neonatal and dirty cells. Expression levels of *Cxcr3* and *Ccr5*, which regulate memory formation and effector versus memory fate decision in CD8+ T cells^96^, were also elevated in VM and neonatal cells,. In contrast, *Ccr9*, involved in T cell development^97^, was more highly expressed in TN and adult cells. Thus, perhaps distinct combinations of cytokine receptors expressed by different naïve subsets set up specific cellular environments and prepare cells for sensing different immune stimuli, for example, to initiate neonatal and VM bystander activation^16,86,88^. *Ccr2*, *Ccr5* and *Cxcr3* were predicted to be activated by *Eomes*, and we observed enhanced binding of *Eomes* at these genes in neonatal cells. *Eomes* may function by regulating these cytokine receptors in naïve cells, helping to preprogram naïve subsets to adopt different activation kinetics and effector potentials upon activation.

## Methods

### Mice

Thy1.1(#000406) and C57Bl/6J (#000664) mice were purchased from Jackson Labs. Heterozygous Lin28b transgenic (Lin28Tg) mice^98^ were provided by Leonid A. Pobezinsky (University of Massachusetts, Amherst, MA) and were maintained as heterozygotes by crossing to C57Bl/6J mice.

### CD8+ sorting for RNA-seq and ATAC-seq

Animals were timestamped^31^ at 7d age and let them age up to 2wk, 8wk or 16wk. Splenic CD8+ T cells were isolated from mice (ZsGreen lox stop lox) by positive CD8a immunomagnetic selection (Miltenyi Biotec). Cells were labeled with antibodies against CD44 (IM7), CD8 (53-6.7), CD4 (GK1.5), and CD49d (R1-2). The cells were then sorted to >95% purity on a FACSAria III (BD Biosciences). VM (CD44hi CD49dlo) and TN (CD44lo CD49dhi) cells were sorted from naive cells. Populations of naive splenocyte CD8+ T cells in Lin28bTg and wildtype (WT) mice were sorted from 8-9 weeks female mice: TN (CD44lo-CD122lo) and VM (CD44hi-CD122hi).

### RNA-seq and ATAC-seq nuclei prep, library preparation and sequencing

For RNA-seq libraries, cells were spun at 1000g for 7’ at 4C and removed the supernatant. Total RNA was isolated with Trizol, with an extra chloroform extraction to remove residual phenol. Stranded RNAseq libraries were prepared with the NEBNext Ultra II Directional RNA kit using 20ng total RNA as input.

For ATAC-seq libraries, cells were spun at 1000g for 7’ at 4C, removed the supernatant and lysed in 1mL trizol. Nuclei were prepared using ‘Buffer H’ for cryopreservation. The nuclei were then thawed and washed in RSB+Tween. The Omni-ATAC protocol^99^ was followed with the following modifications: 25,000 nuclei was used in a 25ul transposition reaction. PCR cycle number per sample was determined based on the original ATAC protocol^40^.

### RNA-seq data analysis

Adaptors were trimmed from raw reads with trim_galore with default parameters. Reads were mapped to the mm10 genome with HISAT2^100^. FeatureCounts^101^ was then used to counts mapped reads for genomic features using GENCODE mouse gene annotation v21 (-s 0 -Q 50). Count matrix of all profiles was normalized and transformed to log2 scale using VST function in DESeq2 (blind = True)^102^. Batch effects across datasets were then removed using R package ComBat^103^, with other cell type factors as covariates. PCA was conducted using prcomp function in R. Correlation of principal components with metadata was performed using eigencorplot function in R package PCAtools^104^. Differential expression between the cell types of interest were performed using DESeq2 with raw counts as inputs, and other factors and dataset sources were used as covariates. GSEA on differential expression results was performed using fgsea^105^ in R (eps = 0.0, minSize = 15, maxSize = 500), with gene sets derived from clusters in Best et al. 2013^106^ describing CD8+ T cell immune response.

### ATAC-seq data analysis

Raw sequencing reads were processed using ENCODE ATAC-seq pipeline v1.1.7. Trimmed adaptor sequences were ‘R1: CTGTCTCTTATACACATCT; R2: CTGTCTCTTATACACATCT’ for profiles in Yee Mon et al, 2021 and Smith et al, 2018. Reads were mapped to the mouse genome (mm10). Paired-end mode was enabled, and peak calling was performed using MACS2^107^ with ‘-f BEDPE’. Irreproducibility Discovery Rate (IDR) analysis ^108^ was enabled for cross-validation to compare replicates. Reads were counted on peaks merged from all samples with featureCounts^101^ (‘-F SAF -s 0 -Q 38’ for profiles in Smith et al, 2018 and ‘-p -F SAF -s 0 -Q 38’ for the rest of profiles).

Count matrix of all profiles were normalized and transformed to log2 scale using VST function in DESeq2^102^ (blind = True). Batch effects across datasets were then removed using R package ComBat^103^, with other cell type factors as covariates. PCA was conducted using prcomp function in R. Correlation of principal components with metadata was performed using eigencorplot function in R package PCAtools^104^. Differential accessibility of peaks between the cell types of interest were performed using DESeq2 with raw counts as inputs, and other factors and dataset sources were used as covariates. Peaks were associated to closest genes using annotatePeaks.pl in HOMER^109^. Motif enrichment analysis was performed using Analysis of Motif Enrichment (AME) in MEME suite^110^, comparing peaks significantly more accessible in one condition to control peaks. Control peaks were randomly selected from peaks that were not differentially accessible in the comparison using fasta-subsample in MEME suite^111^. The number of control peaks selected was mean of the significantly differentially accessible peaks in two conditions in the comparison. JASPAR 2020^112^ core TF motifs in vertebrates was used in the analysis. TF footprint analysis was performed using R package BaGFoot^47^. Samples of each cell type was merged prior to the BaGFoot analysis.

### Network inference

#### Selection of potential target genes

Differential expression analysis was performed between naïve CD8+ T cell subsets for each project. Genes included as potential target genes in the network inference process were the genes differentially expressed with cutoff at adjusted p-value 0.01 in at least one differentially expression comparison for at least one project. In total, 8207 genes were included as potential target genes of the network.

#### Selection of potential regulators

We selected potential regulators by intersecting the mouse TF list constructed by Miraldi et al., 2019 and the differentially expressed genes identified in the previous step. In total, 612 TFs were included as potential regulators of the network.

#### Construction of prior matrices

Peaks called in described ATAC-seq profiles of naïve CD8+ T cell subsets were merged. Position weight matrices of mouse TF motifs were exported from CisBP version 2.00^113^ in Feb 2021. Gene annotation from Gencode vM21 were used to identify genome features. Prior matrices for ATAC-seq were generated based on motif occurrence in the overlapping regions of genome features ±10kb and open chromatin regions, as in Miraldi et al., 2019^24^.

In addition, several TF knockout RNA-seq datasets (Tcf1 from He et al., 2016^84^, Bach2 from Roychoudhuri et al., 2016^41^), TF ChIP-seq datasets (Runx3 from Lotem et al., 2013^114^, Tcf1 from Steinke et al., 2014^115^) and TF network (Bing et al., 2016^84^) in naïve CD8+ T cells were used when constructing the prior matrices. Specifically, for the RNA-seq datasets, differential expression analysis was performed with R package DESeq2 comparing the TF knockout and control samples. Genes with adjusted p-values smaller than 0.05 were classified as differential expression genes, which were included as targets of the TF in the prior matrix. For the TF ChIP-seq datasets, peaks were called using macs2 callpeak^107^, with p-value cutoff at 0.05. To identify potential targets of the TF, motif search was performed in the overlapping regions of genome features ±10kb and ChIP-seq peaks. Genes with motif identified in regions closeby were included as targets of the TF in the prior matrix.

#### Inference framework and final network

The Inferelator framework was as described in Miraldi et al., 2019, with moderate prior reinforcement, model size 10, combined TF activity options. The framework was run five times with random seeds, and regulations appeared in more than three networks were included for the final network.

#### Core TFs for each naïve CD8+ T cell subset

Differential expression analysis comparing VM vs TN, neonatal vs adult, dirty vs clean subsets of naïve CD8+ T cells was conducted using R package DESeq2. Datasets containing the relevant subsets for each comparison were used, with sources of profiles included as covariates. GSEA were conducted based on the differential expression results using predicted targets of each TF in the network. Core TFs for each subset were identified through an approach adapted from Miraldi et al: core TFs were defined as those TFs whose positive targets were enriched on the side of subset of interest, or the TFs whose negative targets were enriched on the side of its counterpart. TF-TF modules with shared targets were identified using the predicted network as described in Miraldi et al., 2019.

### Micro-C library preparation, sequencing and data analysis

Adult CD8+ T cells were magnetically enriched from the spleen. The cells were then FACS sorted on live CD49d-CD8+ CD4-CD44+ CD122+ (For VM) and live CD49d-CD8+ CD4-CD44-CD122-(For TN). Micro-C libraries were made using Dovetail Genomics Micro-C Kit (v 1.0). Sorted cells were centrifuged at 500 xg for 5 minutes at 4°C to remove sheath buffer. After removing the supernatant, cells were resuspended in 1 ml of 1x PBS and split into aliquots of 500,000 cells or less. Cells were spun again at 500xg for 5 minutes at 4°C, the supernatant was removed and resuspended in 1 ml of 1x PBS and mixed gently, counted, and spun hard at 3,000 xg for 5 minutes at 4°C. Supernatant was removed, and the pellet was stored “dry” at −80°C. Cells were removed after an overnight and thawed rapidly at room temperature for processing. Briefly, cells were resuspended in 1ml of 1x PBS and crosslinked with 0.3M DSG and 1% formaldehyde. After washing the cells, they were digested with varying amounts of MNase (based on cell counts) in nuclease digest buffer with MgCl_2_. Digestion was stopped by the addition of 0.5M EGTA and cells lysed by the addition of 20% SDS. Lysate QC was performed using an Agilent 5200 Fragment analyzer. Chromatin bound DNA ends were repaired and A-tailed using the End Polishing enzyme mix and buffer. Dovetail bridge was then ligated to the chromatin bound DNA free ends following the Micro-C kit protocol then subjected to intra-aggregate ligation to create chromatin-chromatin long range interaction. Finally, the DNA bound to chromatin was recovered by proteinase K treatment and crosslink reversal and purified using SPRIselect beads (Beckman Coulter). The NEB NEBNext Ultra II DNA Library Prep Kit was used for Illumina sequencing. Final libraries were size selected to be 300 – 1000 bp using the Pippin (Sage Science). Libraries were sequenced initially on one HiSeq lane, the library pool was rebalanced, checked using a MiSeq lane, and finally sequenced deeply on a NovaSeq S4 lane. Reads were mapped to mouse genome (mm10) with bwa mem (−5SP -T0 -R “@RG\tID:$rg\tSM:$rg\tLB:$rg\tPL:ILLUMINA\tPU:none”). The alignments were converted into .pairs, sorted and deduplicated by pairtools. Data from all samples were merged by pairtools and converted to .hic file by Juicer v1.6.2^116^.

### Network validation with the Micro-C dataset

Chromatin loops were found with HiCCUPS (juicer_tools_1.22.01, --cpu -r 1000,5000,10000,25000,50000 -k KR -f 0.001,0.001,0.001,0.001,0.001 --ignore-sparsity) from the .hic file. “Open” loops were found by overlapping the loop ends with ATAC-seq peaks. Each end of the open loops were associated with genes within 10kb with bedmap in BEDOPS^117^.

Scanning of TF motifs in the open loop ends was performed with R package motifmatchr^118^. The set of Micro-C derived regulations were then derived by linking the TFs with motif occurrences to genes associated to either end of the loops. 10000 random networks were generated by permutating the target genes in the network for 10000 times.

### Network validation with TF perturbation RNA-seq profiles

*Eomes*-overexpression (OE) RNA-seq dataset from CD8+ single positive cells were generated in Istaces et al. 2019^51^. Log2 fold changes between wildtype samples and *Eomes* overexpressed samples were selected for predicted *Eomes* targets and background genes (other potential target genes during network inference). Distributions of log2 fold changes of *Eomes* positive/negative targets and background genes were compared with Wilcoxon test.

*Foxo1* knockout RNA-seq profiles of naïve CD8+ T cells were generated in Ouyang et al. 2009^53^. Log2 fold changes between the *Foxo1* knockout sample and the wildtype sample were selected for predicted *Foxo1* targets and background genes. Distributions of log2 fold changes of *Foxo1* positive/negative targets and background genes were compared with Wilcoxon test.

### CUT&Tag library preparation and sequencing

#### Transposome preparation

CUT&Tag was performed using novel a fusion construct pAGLL developed in the Genomics Innovation Hub at Cornell University, which consists of a protein-A/G fusion to Tn5, separated by a long flexible linker. The starting vector (pAG-Tn5) was generated by GeneArt Custom Gene Synthesis (Thermo Fisher Scientific). To improve the functional range of the pAG-Tn5 fusion, the vector was then modified using rounds of site-directed mutagenesis (Q5 Site-directed mutagenesis kit #E0554S, NEB) to first introduce an M56L mutation in Tn5 (Primers:Tn5M56L_F: 5’-TAGCAAAGCATTGCAAGAAGGTG,Tn5M56L_R: 5’-CCTTCGCTTGAAATGGTAATG) to prevent the expression of the Tn5 inhibitor protein (Inh)^119^. The second round of mutagenesis increased the number of GGGS repeats to 7, as well as add an additional peptide linker of GEGQGQGQGPG^120^ using primers: pAGTn5_linkinsF: 5’-GGTAGCGGTGAAGGCCAGGGCCAAGGCCAGGGGCCGGGACATATGATTACCAGCGCACTGCATCG and pAGTn5_linkins_R: 5’-ACCGCCACCGCTACCACCGCCACCGCTACCACCGCCACCGCTACCGCCACCGCCAGA. The resulting linker sequence in the pAGLL construct is: DDDKEFGGGGSGGGGSGGGGSGGGGSGGGGSGGGGSGGGGSGEGQGQGQGPGH. After mutagenesis, all plasmids were sequenced verified (Plasmidsaurus).

The pAGLL plasmid was transformed into *E.coli* (C1301, NEB) and the fusion proteins were purified via affinity chromatography as described^121^. Briefly, a 1L culture was inoculated with an overnight starter culture (1:50 dilution) in 2XYT media with carbencillin and grown to a OD600 of 0.6. The culture was chilled on ice for 10 minutes prior to induction with 1mM IPTG. The culture was then grown for an additional 4-6 hours at room temperature. The culture was centrifuged and the bacterial pellets washed in cold HEGX buffer (20mM HEPES at pH 7.3, 800mM NaCl, 1mM EDTA, 10% glycerol, 0.2% Triton X-100). The bacterial pellets were frozen and stored at −80°C. Pellets were resuspended in 50mL cold HEGX buffer plus protease inhibitors and sonicated using a US SOLID sonicator set to 50% power for 4 minutes using a 5sec on/5sec off program and waiting 1 minute between rounds of sonication. After centrifugation to remove insoluble material, genomic DNA was precipitated by adding 2.5% of lysate volume with 10% PEI. The cleared lysate was then added to a chitin resin (NEB) column and allowed to bind. After multiple washes with HEGX, bound protein was eluted using HEGX with 50mM DTT and sealing the column for 48 hours. The elution was then collected, dialyzed (100mM HEPES at pH 7.3, 200mM NaCl, 0.2mM EDTA, 20% glycerol, 0.2% Triton X-100, 2mM DTT) and the functional concentration of pAGLL determined using qPCR. Purified protein was then aliquoted and stored at −80°C.

To generate the pAGLL adapter transposome, ME-A (5′-TCGTCGGCAGCGTCAGATGTGTATAAGAGACAG-3′) and ME-B (5′-GTCTCGTGGGCTCGGAGATGTGTATAAGAGACAG-3′.) were annealed to ME-Rev(5’PHOS/CTGTCTCTTATACACATCT) separately. Then equimolar mixture of preannealed ME-A and ME-B oligonucleotides were used to load pAGLL for 21 hours at 25°C with 600rpm shaking in an Eppendorf Thermomixer.

#### CUT&Tag Library Preparation

Spleens were harvested from mice and mashed through 40um strainer. Briefly, 250,000 mouse CD8+ cells were magnetically isolated using CD8a microbeads (Miltenyi Biotec, 130-117-044) from mouse spleens, fixed with 1% formaldehyde (VWR, 15710) at room temperature (RT) for 10 minutes, and then slowly frozen in PBS containing 10% DMSO. The CUT&Tag protocol was adapted from Kaya-Okur et al with modifications^29^. For the CUT&Tag libraries preparation, frozen cells were thawed and immobilized onto concanavalin A-coated magnetic beads (Bangs Laboratories, BP531). The bead-bound cells were resuspended in antibody binding buffer with BSA (20mM HEPES at pH 7.5, 150mM NaCl, 0.5mM spermidine, 0.01% digitonin, 2mM EDTA, 1% nuclease-free BSA) and a 1:50 dilution of the appropriate primary antibody (anti-Eomes, Abcam, ab23345 and rabbit IgG, Cell Signaling Technology, 66362). Primary antibody incubation was performed on a nutator platform overnight at 4°C. Secondary antibody binding was omitted. The cells were then washed twice in dig-wash buffer (20 mM HEPES at pH 7.5, 150 mM NaCl, 0.5 mM spermidine, 0.01% digitonin, complete Protease Inhibitor EDTA-Free) with 5 minutes nutation. Next, the cells were resuspended in dig-300 buffer (20 mM HEPES at pH 7.5, 300 mM NaCl, 0.5 mM spermidine, 0.01% digitonin, complete Protease Inhibitor EDTA-Free) containing 1 μM homemade pAGLL-Tn5 adapter complex and incubated for 1 hour at RT on a nutator. The cells were washed twice with dig-300 buffer (20 mM HEPES at pH 7.5, 300 mM NaCl, 0.5 mM spermidine, 0.01% digitonin, complete Protease Inhibitor EDTA-Free) with 5 minutes nutation and then resuspended in tagmentation buffer (1x rCutSmart Buffer NEB #B6004S, 10% dimethylformamide, 150 mM NaCl). Tagmentation was carried out by incubating the cells for 1 hour at 37°C with 500rpm shaking in an Eppendorf Thermomixer. To stop the tagmentation and reverse the crosslinks, 50 µL reverse-crosslinking buffer (3.33 µL 0.5M EDTA, 1 µL 10% SDS, 2 µL Proteinase K NEB #P8111S, 43.67 µL EB buffer) was added to each sample. After incubating on a thermomixer for 30 minutes at 37°C and 10 minutes at 55°C, the DNA was purified using the Zymo DNA Clean & Concentrator-5 kit. The libraries were PCR amplified using unique dual UDI Nextera primers and 2X NEBNext High Fidelity master mix (NEB# M0541) using the program: 72°C 5min, 98°C 45 sec, followed by 16 cycles of 98°C 10sec, 63°C 15 sec, final 72°C 5 min. PCR products were purified using SparQ beads (Quanta Bio) and resuspended in 0.1X TE buffer. Libraries were then checked on a Fragment analyzer (Agilent) and size selected using a Pippin HT (Sage Sciences). Libraries were sequenced on an Illumina NextSeq 500 platform using 2×150 bp paired-end sequencing.

### CUT&Tag data analysis

Trim Galore v0.6.5^122,123^ was used for quality and adaptor trimming (--quality 20 --length 20 --paired). Bowtie2 v2.5.1^124^ was used to map reads on mouse genome (mm10). Reads mapped to mitochondria and blacklist regions^125^ were removed. Picard v2.26.1^126^ was to mark and remove duplicated reads (VALIDATION_STRINGENCY=LENIENT). bamCoverage v3.5.5^127^ was used to generate bigwig files (--normalizeUsing CPM --binSize 50 -effectiveGenomeSize 2,652,783,500). Macs3 v3.0.0^107^ was used to call peaks for Eomes adult and lin28Tg samples, with all IgG adult and lin28Tg samples as control (-f BAMPE -g mm -q 0.01 --fe-cutoff 10 --nolambda). Peaks were further filtered at q-value 1e-20, 1e-50, 1e-100, 1e-150, 1e-180. Default parameters were used unless described otherwise.

To call differential peaks, macs3 callpeak was first ran for adult and Lin28Tg samples separately, with IgG samples from each sample type as controls (-B -f BAMPE -g mm --nolambda --keep-dup all). Macs3 bdgdiff was then used to identify differential peaks between adult and Lin28Tg samples (--d1 3146753 --d2 1931794, which were the actual effective depths of adult and Lin28Tg samples obtained from the previous callpeak step).

R package monaLisa v1.2.0^128^ was used to perform motif enrichment analysis on CUT&Tag peaks (filtered for peaks with q-value less than 1e-150) with background sampled from ATAC-seq peaks. CUT&Tag peaks were associated to nearest genes using annotatePeaks.pl from Homer^109^. Overrepresentation analysis based on hypergeometric test was performed with R package fgsea v1.22.0^105^ using gene sets from Hallmark^129^, Reactome^130^ and those derived from clusters in Best et al. 2013^106^ describing CD8+ T cell immune response. Fisher’s exact test was used to evaluate the enrichment of CUT&Tag-derived TF targets and those from the predicted network.

## Data availability

Raw and analyzed data generated in this paper were deposited to GEO under accessions GSE276151 (Micro-C), GSE276148 (CUT&Tag), GSE276149 (RNA-seq), GSE276147 (ATAC-seq) and GSE276146 (RNA-seq).

## Code availability

All code for analysis is available upon request.

## Acknowledgments

We thank the Cornell Biotechnology Resource Center (BRC) Genomics, Flow Cytometry, and Transcriptional Regulation and Gene Expression (RRID: SCR_021727; SCR_021740; SCR_022532) facilities for support.

## Author Contributions

Conceptualization H.Z. and A.G.; Experiments Y.J., A.J.M., E.A.F., C.T., R.K.P. and N.L.S.; Data analysis H.Z., J.D.C. and P.R.M.; Formal analysis and visualization H.Z.; Supervision A.G., B.D.R. and J.K.G.; Writing H.Z. and A.G.

