## Supplementary Figures for "Deciphering gene regulatory programs underlying functionally divergent naïve T cell subsets"

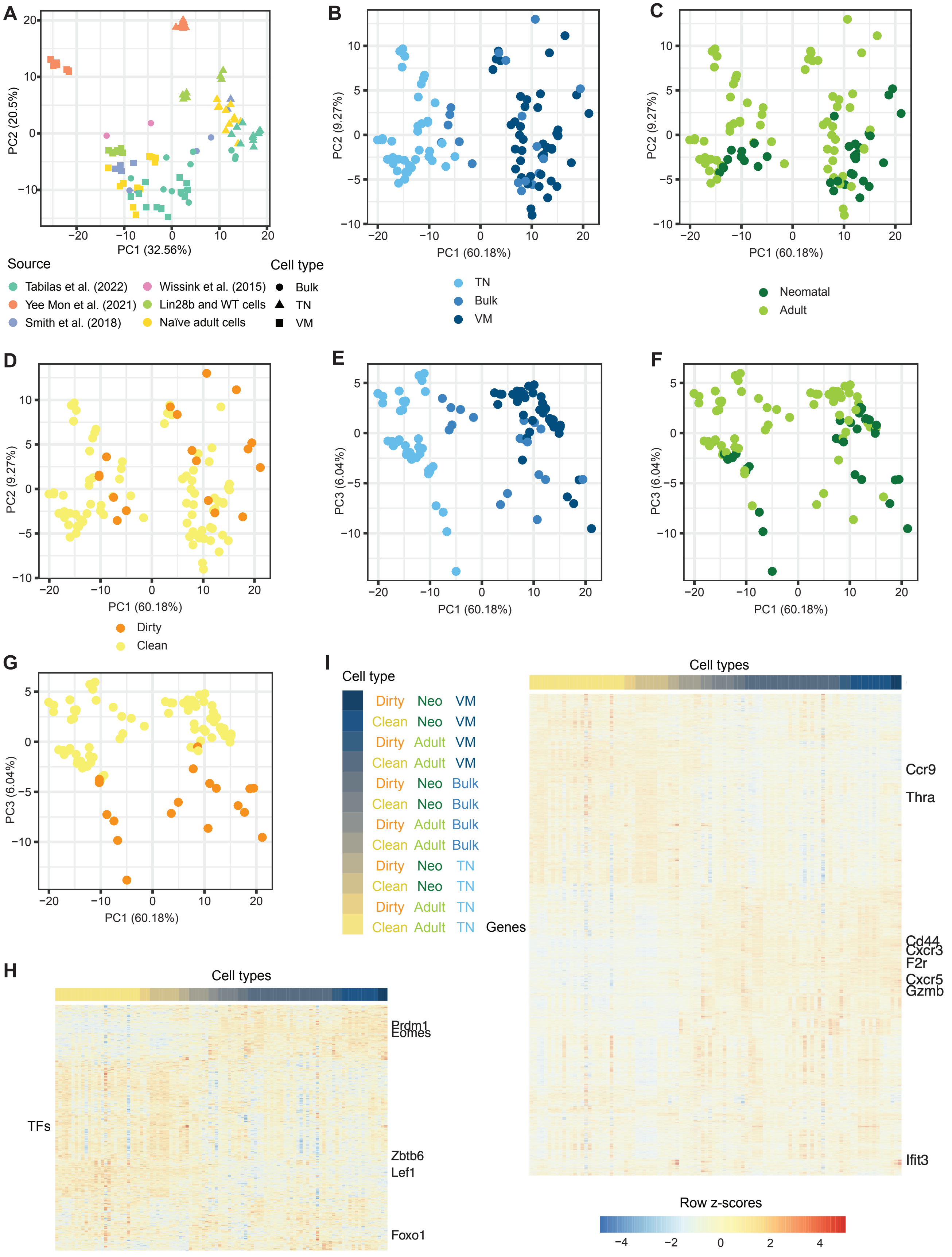
A
PC1 (32.56%)
PC2 (20.5%)
Source
Tabilas et al. (2022)
Yee Mon et al. (2021)
Smith et al. (2018)
Wissink et al. (2015)
Lin28b and WT cells
Naïve adult cells
Cell type
Bulk
TN
VM
B
PC1 (60.18%)
PC2 (9.27%)
TN
Bulk
VM
C
Neomatal
Adult
D
Dirty
Clean
E
PC3 (6.04%)
F
G
H
Cell types
TFs
Prdm1
Eomes
Zbtb6
Lef1
Foxo1
I
Cell type
Dirty Neo VM
Clean Neo VM
Dirty Adult VM
Clean Adult VM
Dirty Neo Bulk
Clean Neo Bulk
Dirty Adult Bulk
Clean Adult Bulk
Dirty Neo TN
Clean Neo TN
Dirty Adult TN
Clean Adult TN
Genes
Cell types
Ccr9
Thra
Cd44
Cxcr3
F2r
Cxcr5
Gzmb
Ifit3
Row z-scores

**Supplementary Fig. 1: Transcriptional landscapes of naïve CD8+ T cell subsets**

(A) PCA of RNA-seq profiles from naïve CD8+ T cells subsets, before batch effect removal. (B-D) PCA of RNA-seq profiles from naïve CD8+ T cells subsets colored by cell types (B), age (C) and microbial environment (D), showing PC1 and 2. (E-G) PCA of RNA-seq profiles from naïve CD8+ T cells subsets colored by cell types (E), age (F) and microbial environment (G), showing PC1 and 3. (H) Heatmap showing TF expression levels across naïve CD8+ T cell subsets, scaled by rows (samples). (I) Heatmap showing gene expression levels across naïve CD8+ T cell subsets, scaled by rows (samples). Related to Fig. 1.

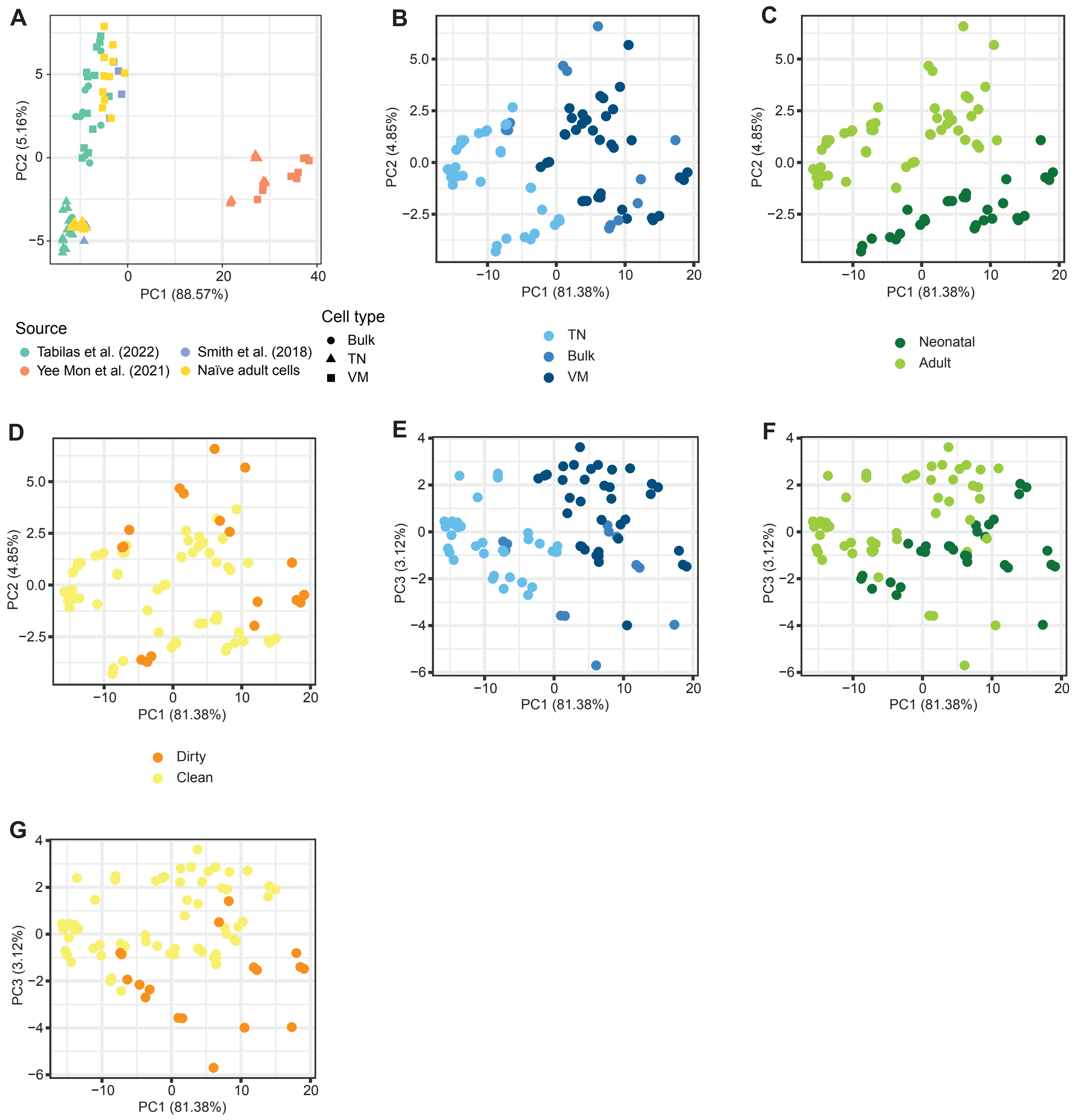


**Supplementary Fig. 2: Naïve CD8+ T cell subsets have diverse chromatin accessibility and TF activity patterns**

(A) PCA of ATAC-seq profiles from naïve CD8+ T cells subsets, before batch effect removal. (B-D) PCA of ATAC-seq profiles from naïve CD8+ T cells subsets colored by cell types (B), age (C) and microbial environment (D), showing PC1 and 2. (E-G) PCA of ATAC-seq profiles from naïve CD8+ T cells subsets colored by cell types (E), age (F) and microbial environment (G), showing PC1 and 3. Related to Fig. 2.

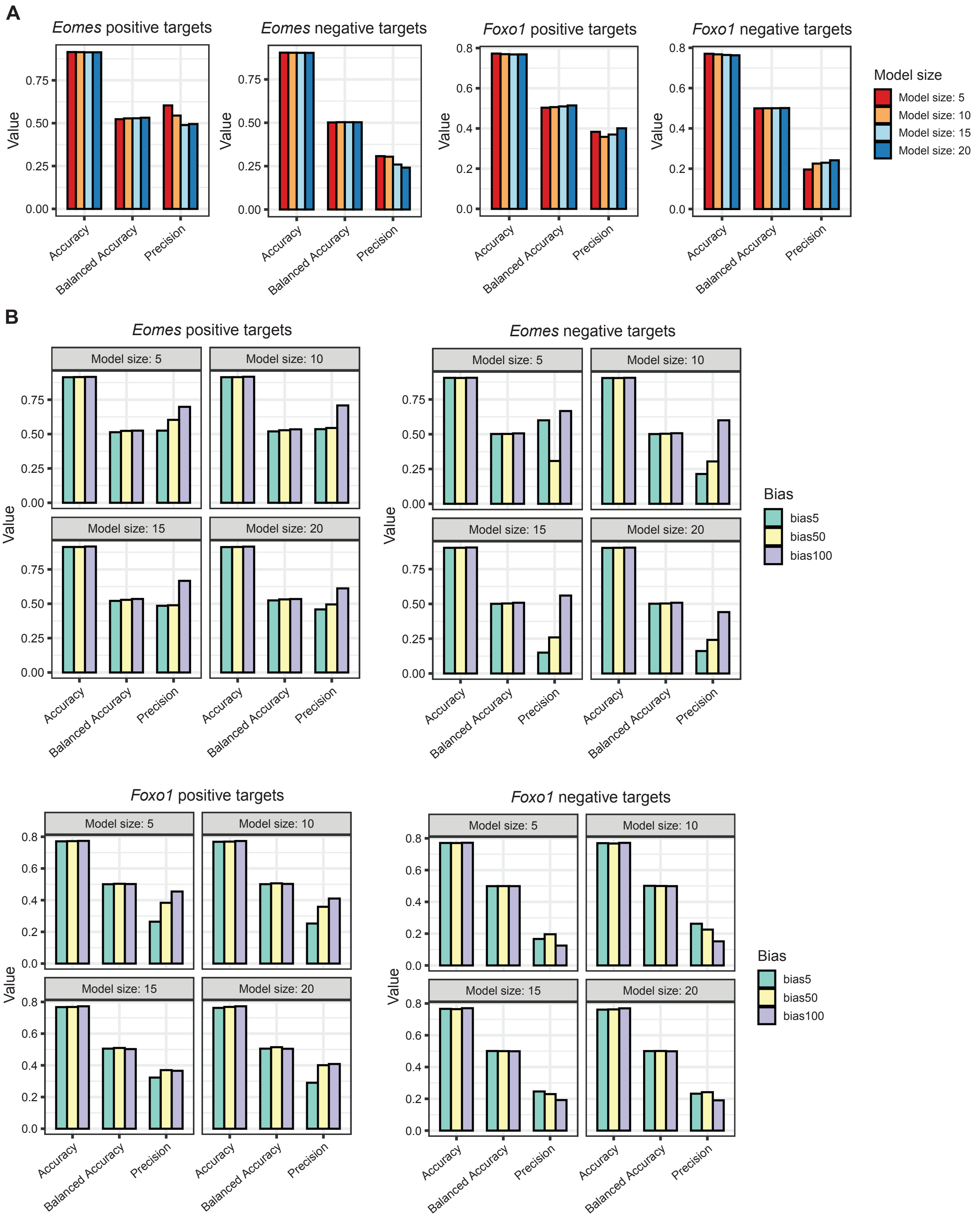

A
Eomes positive targets
Eomes negative targets
Foxo1 positive targets
Foxo1 negative targets
Value
Accuracy
Balanced Accuracy
Precision
Model size
Model size: 5
Model size: 10
Model size: 15
Model size: 20
B
Eomes positive targets
Eomes negative targets
Foxo1 positive targets
Foxo1 negative targets
Model size: 5
Model size: 10
Model size: 15
Model size: 20
Value
Accuracy
Balanced Accuracy
Precision
Bias
bias5
bias50
bias100

C

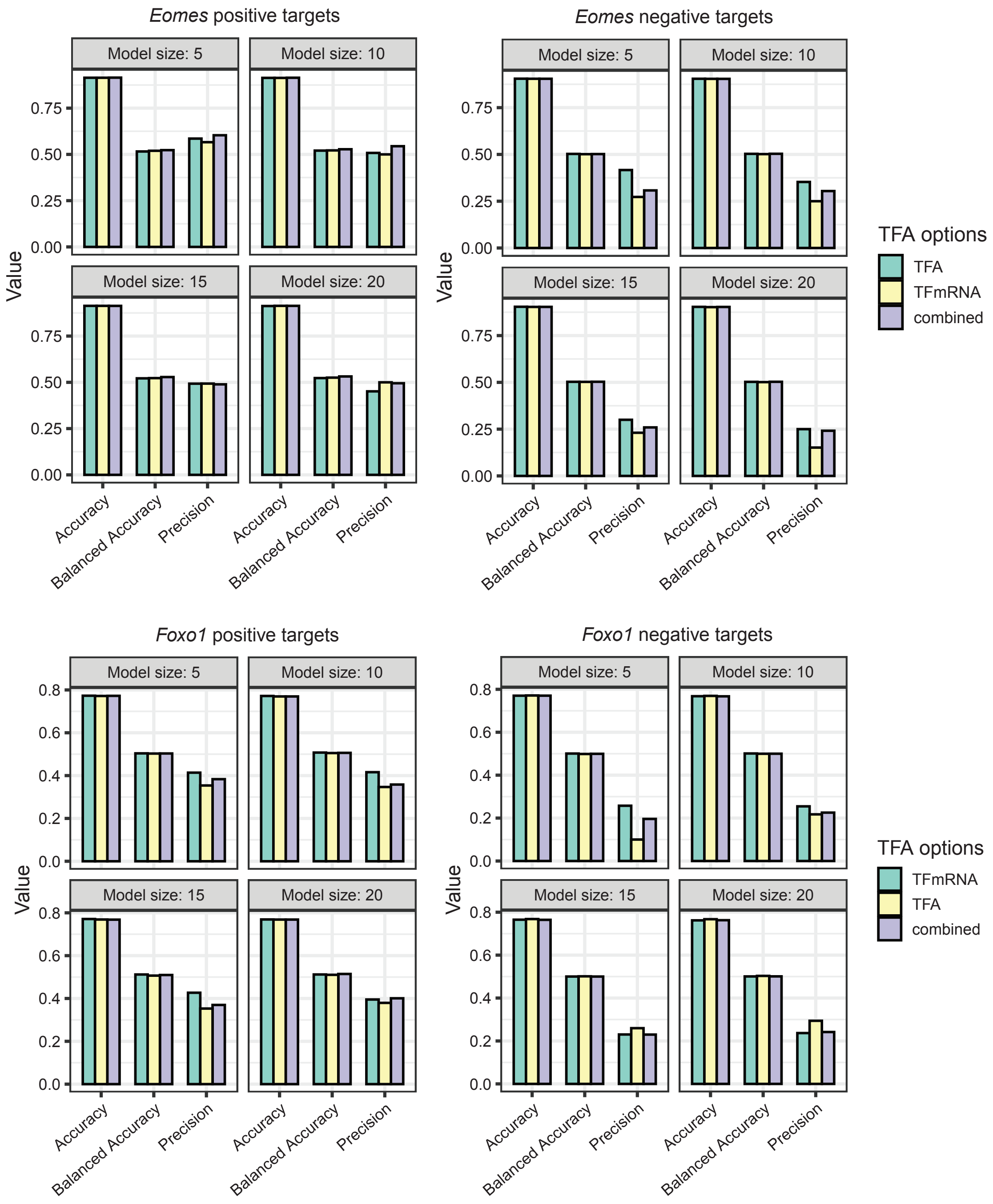
Eomes positive targets
Eomes negative targets
Foxo1 positive targets
Foxo1 negative targets
Model size: 5
Model size: 10
Model size: 15
Model size: 20
Value
Accuracy
Balanced Accuracy
Precision
TFA options
TFA
TFmRNA
combined

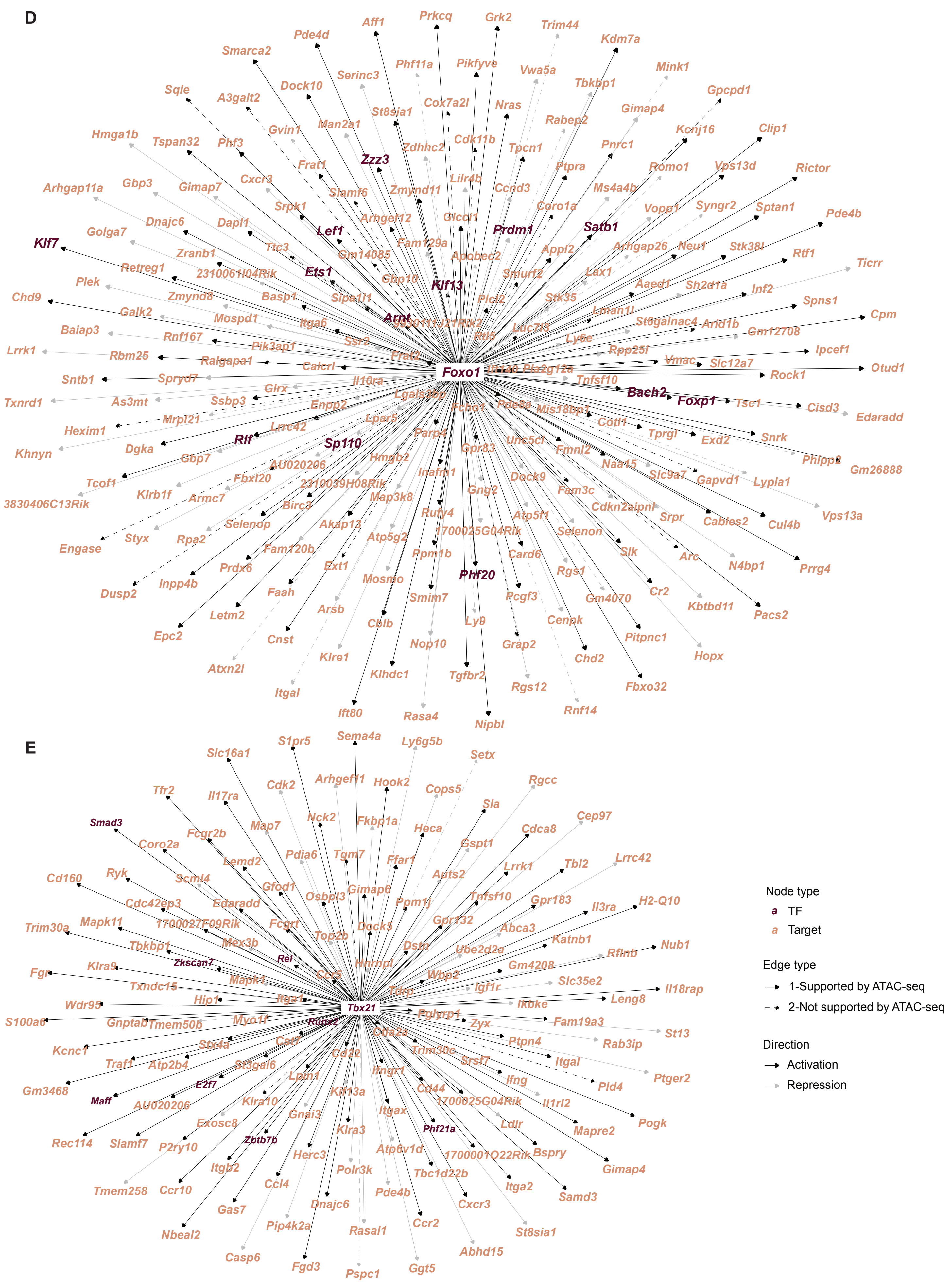
D
Foxo1
Zzz3
Lef1
Prdm1
Satb1
Klf7
Ets1
Klf13
Arnt
Bach2
Foxp1
Rlf
Sp110
Phf20
Aff1
Prkcq
Grk2
Trim44
Kdm7a
Pde4d
Smarca2
Phf11a
Pikfyve
Vwa5a
Mink1
Serinc3
Dock10
Sqle
A3galt2
St8sia1
Cox7a2l
Nras
Tbkbp1
Gpcpd1
Gimap4
Man2a1
Gvin1
Rabep2
Kcnj16
Clip1
Hmga1b
Tspan32
Phf3
Zdhhc2
Cdk11b
Tpcn1
Pnrc1
Frat1
Romo1
Vps13d
Rictor
Lilr4b
Cxcr3
Ptpra
Arhgap11a
Gbp3
Gimap7
Slamf6
Zmynd11
Ccnd3
Ms4a4b
Srpk1
Coro1a
Vopp1
Syngr2
Sptan1
Pde4b
Arhgef12
Glcci1
Dnajc6
Dapl1
Golga7
Fam129a
Appl2
Arhgap26
Neu1
Stk38l
Rtf1
Ticrr
Ttc3
Zranb1
Gm14085
Apobec2
Retreg1
2310061I04Rik
Smurf2
Lax1
Aaed1
Sh2d1a
Inf2
Plek
Gbp10
Spns1
Zmynd8
Basp1
Sipa1l1
Plcl2
Stk35
Chd9
Lman1l
Cpm
Galk2
Itga6
St6galnac4
Arid1b
Mospd1
Baiap3
Ssr2
Luc7l3
Ly6e
Gm12708
Rnf167
Pik3ap1
Rpp25l
Ipcef1
Lrrk1
Rbm25
Frat2
Vmac
Slc12a7
Otud1
Ralgapa1
Calcrl
Pla2g12a
Rock1
Sntb1
Spryd7
Il10ra
Tnfsf10
Glrx
Txnrd1
As3mt
Ssbp3
Lgals3bp
Tsc1
Cisd3
Edaradd
Enpp2
Mis18bp1
Hexim1
Mrpl21
Lrrc42
Lpar5
Cotl1
Tprgl
Exd2
Snrk
Khnyn
Dgka
Parp4
Unc5cl
Fmnl2
Phlpp2
Gm26888
Gbp7
AU020206
Hmgb2
Gpr83
Naa15
Slc9a7
Gapvd1
Lypla1
Tcof1
Fbxl20
Inafm1
Dock9
Klrb1f
2310039H08Rik
Gng2
Fam3c
Vps13a
3830406C13Rik
Armc7
Birc3
Map3k8
Cdkn2aipnl
Srpr
Cables2
Cul4b
Engase
Styx
Rpa2
Selenop
Rufy4
Atp5f1
Akap13
1700025G04Rik
Selenon
Arc
N4bp1
Prrg4
Atp5g2
Fam120b
Ppm1b
Card6
Slk
Prdx6
Ext1
Rgs1
Dusp2
Inpp4b
Mosmo
Cr2
Faah
Smim7
Pcgf3
Gm4070
Kbtbd11
Pacs2
Letm2
Arsb
Epc2
Cnst
Cblb
Ly9
Cenpk
Pitpnc1
Nop10
Grap2
Hopx
Klre1
Chd2
Atxn2l
Klhdc1
Tgfbr2
Rgs12
Fbxo32
Itgal
Ift80
Rasa4
Rnf14
Nipbl
E
Tbx21
Smad3
Zkscan7
Rel
Runx2
E2f7
Maff
Zbtb7b
Phf21a
S1pr5
Sema4a
Ly6g5b
Setx
Slc16a1
Cdk2
Arhgef11
Hook2
Cops5
Rgcc
Tfr2
Il17ra
Sla
Nck2
Fkbp1a
Cep97
Map7
Heca
Fcgr2b
Coro2a
Gspt1
Cdca8
Pdia6
Tgm7
Lemd2
Ffar1
Lrrk1
Tbl2
Lrrc42
Cd160
Ryk
Scml4
Auts2
Gfod1
Gimap6
Osbpl3
Tnfsf10
Gpr183
Il3ra
H2-Q10
Edaradd
Ppm1j
Cdc42ep3
Mapk11
1700027F09Rik
Fcgrt
Gpr132
Abca3
Trim30a
Dock5
Katnb1
Mex3b
Top2b
Dstn
Nub1
Tbkbp1
Ube2d2a
Rflnb
Klra9
Hnrnpl
Gm4208
Fgr
Ccr5
Wbp2
Slc35e2
Txndc15
Mapk1
Igf1r
Il18rap
Hip1
Wdr95
Itga1
Ikbke
Leng8
S100a6
Gnptab
Tmem50b
Myo1f
Pglyrp1
Zyx
Fam19a3
St13
Rab3ip
Ptpn4
Kcnc1
Stx4a
Ctla2a
Trim30c
Traf1
Atp2b4
Csf1
Cd82
Srsf7
Itgal
Ptger2
St3gal6
Lpin1
Ifngr1
Ifng
Pld4
Gm3468
AU020206
Klra10
Kif13a
Cd44
Il1rl2
1700025G04Rik
Exosc8
Gnai3
Itgax
Mapre2
Pogk
Ldlr
Rec114
Slamf7
P2ry10
Klra3
Bspry
Herc3
Atp6v1d
1700001O22Rik
Gimap4
Itgb2
Polr3k
Tbc1d22b
Itga2
Samd3
Tmem258
Ccr10
Ccl4
Pde4b
Gas7
Dnajc6
Cxcr3
Pip4k2a
Rasal1
Ccr2
St8sia1
Nbeal2
Abhd15
Casp6
Fgd3
Pspc1
Ggt5
Node type
a TF
a Target
Edge type
1-Supported by ATAC-seq
2-Not supported by ATAC-seq
Direction
Activation
Repression

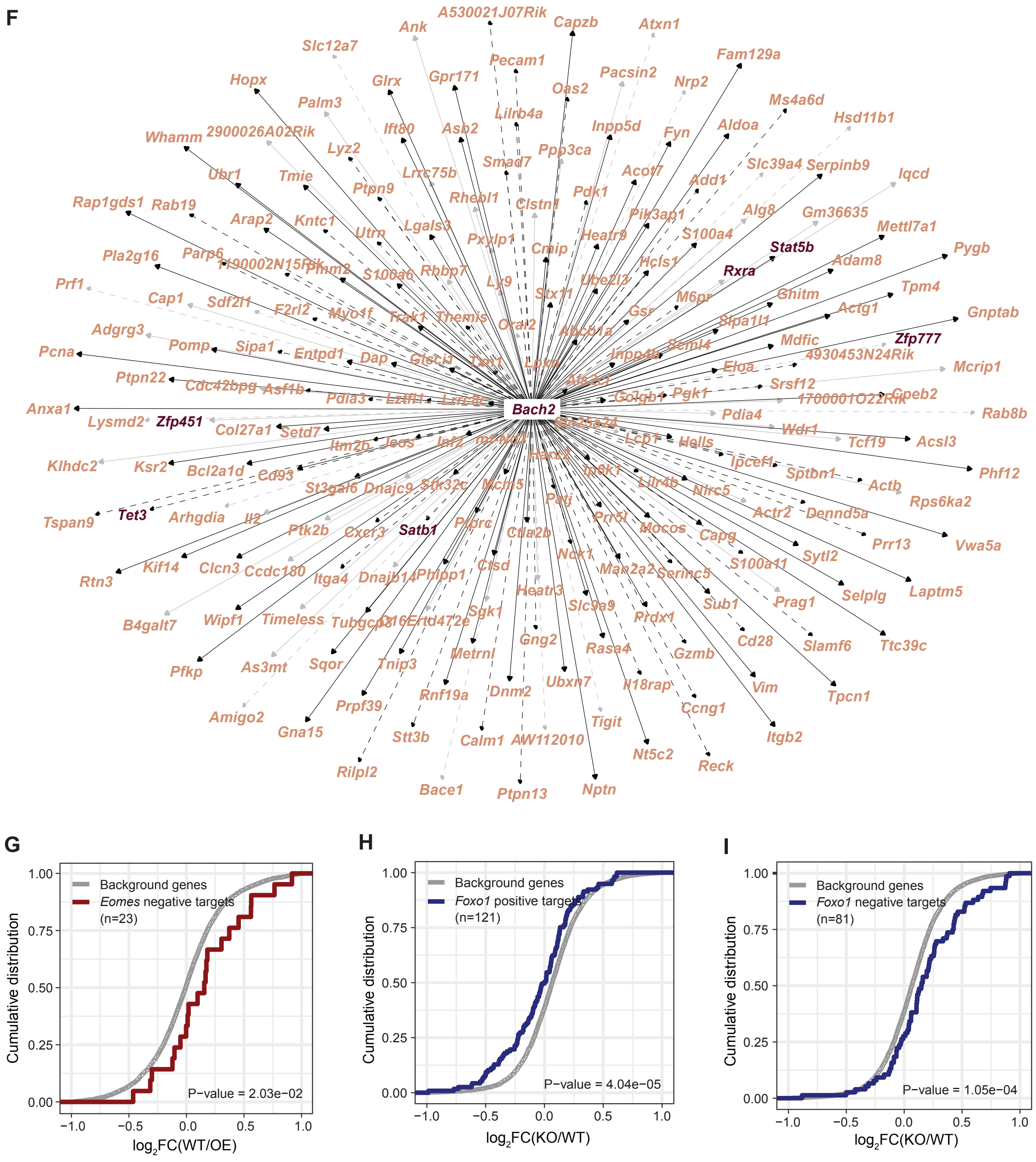


**Supplementary Fig. 3: A transcriptional regulatory network for naïve CD8+ T cells**

(A-C) Evaluations on the predicted Eomes and Foxo1 targets across parameters model size (A), bias (prior reinforcement, B), TF activity (TFA) options (C). (D-F) Predicted targets of *Foxo1* (D), *Tbx21* (E) and *Bach2* (F) in the network. (G) Cumulative distribution plot showing significant differences in distributions of $log_2FC$ (WT/Eomes OE; x-axis) between predicted negative targets of Eomes and background genes (Wilcoxon test). (H-I) Cumulative distribution plot showing significant differences in distributions of $log_2FC$ (Foxo1 KO/WT; x-axis) between predicted positive (H) and negative (I) targets of Foxo1 and background genes (Wilcoxon test). Related to Fig. 3.

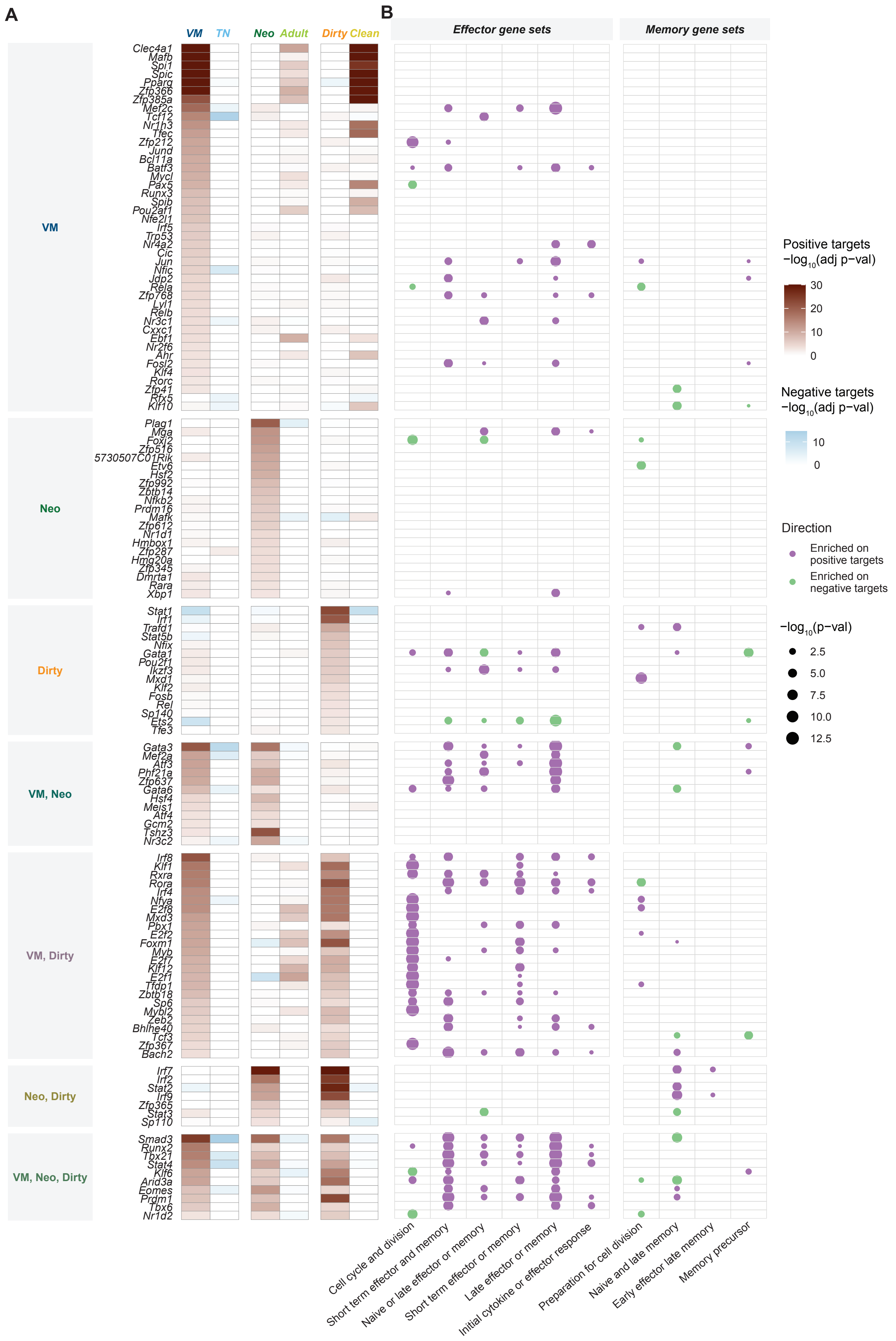
A
B
VM
TN
Neo
Adult
Dirty
Clean
Effector gene sets
Memory gene sets
VM
Clec4a1
Mafb
Spi1
Spic
Pparg
Zfp366
Zfp385a
Mef2c
Tcf12
Nr1h3
Tfec
Zfp212
Jund
Bcl11a
Batf3
Mycl
Pax5
Runx3
Spib
Pou2af1
Nfe2l1
Irf5
Trp53
Nr4a2
Cic
Jun
Nfic
Jdp2
Rela
Zfp768
Lyl1
Relb
Nr3c1
Cxxc1
Ebf1
Nr2f6
Ahr
Fosl2
Klf4
Rorc
Zfp41
Rfx5
Klf10
Neo
Plag1
Mga
Foxj2
Zfp516
5730507C01Rik
Etv6
Hsf2
Zfp992
Zbtb14
Nfkb2
Prdm16
Mafk
Zfp612
Nr1d1
Hmbox1
Zfp287
Hmg20a
Zfp345
Dmrta1
Rara
Xbp1
Dirty
Stat1
Irf1
Trafd1
Stat5b
Nfix
Gata1
Pou2f1
Ikzf3
Mxd1
Klf2
Fosb
Rel
Sp140
Ets2
Tfe3
VM, Neo
Gata3
Mef2a
Atf3
Phf21a
Zfp637
Gata6
Hsf4
Meis1
Atf4
Gcm2
Tshz3
Nr3c2
VM, Dirty
Irf8
Klf1
Rxra
Rora
Irf4
Nfya
E2f8
Mxd3
Pbx1
E2f2
Foxm1
Myb
E2f7
Klf12
E2f1
Tfdp1
Zbtb18
Sp6
Mybl2
Zeb2
Bhlhe40
Tcf3
Zfp367
Bach2
Neo, Dirty
Irf7
Irf2
Stat2
Irf9
Zfp365
Stat3
Sp110
VM, Neo, Dirty
Smad3
Runx2
Tbx21
Stat4
Klf6
Arid3a
Eomes
Prdm1
Tbx6
Nr1d2
Cell cycle and division
Short term effector and memory
Naive or late effector or memory
Short term effector or memory
Late effector or memory
Initial cytokine or effector response
Preparation for cell division
Naive and late memory
Early effector late memory
Memory precursor
Positive targets
−log10(adj p−val)
30
20
10
0
Negative targets
−log10(adj p−val)
10
0
Direction
Enriched on positive targets
Enriched on negative targets
−log10(p−val)
2.5
5.0
7.5
10.0
12.5

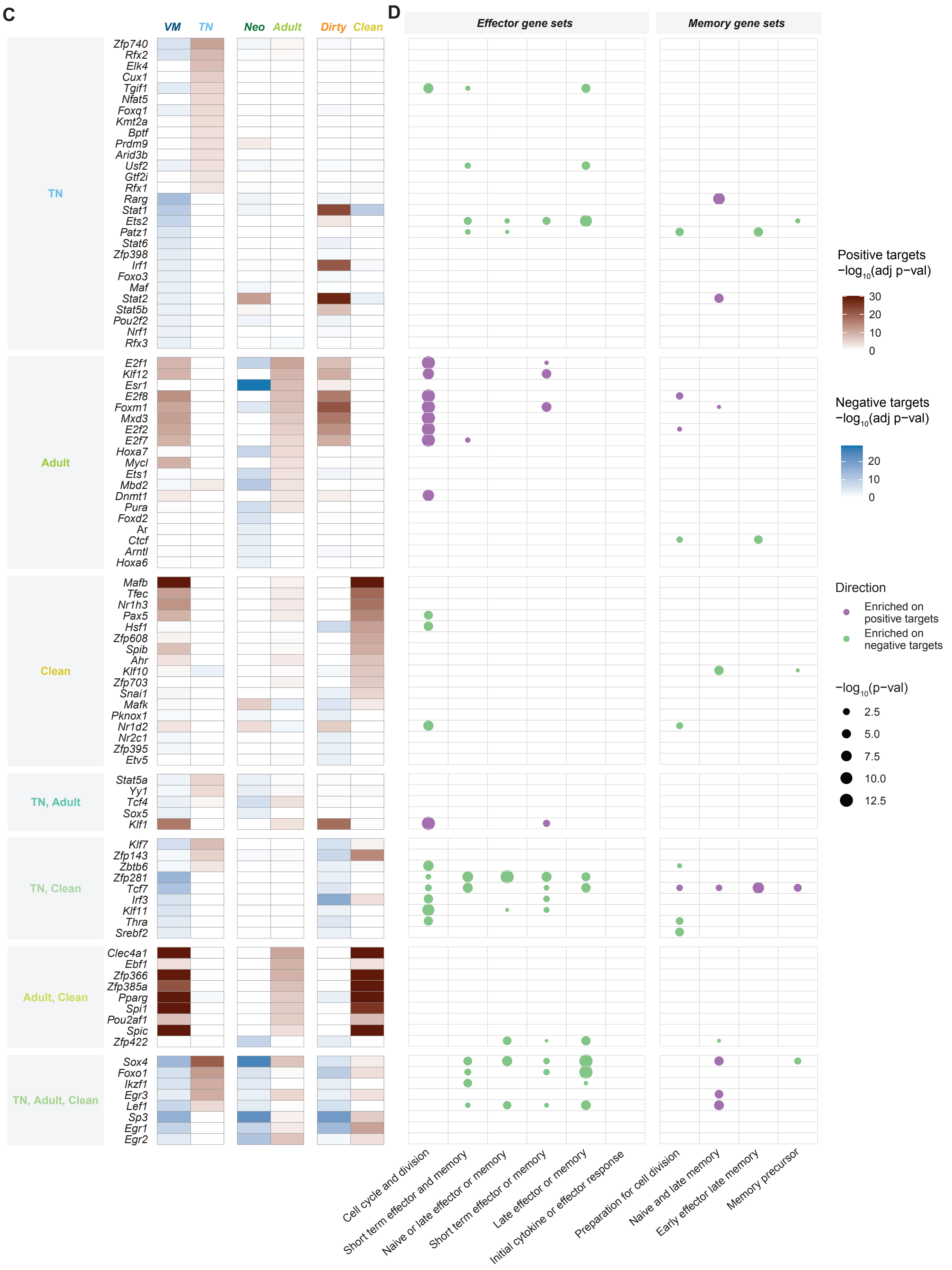

C
D
VM
TN
Neo
Adult
Dirty
Clean
Effector gene sets
Memory gene sets
TN
Zfp740
Rfx2
Elk4
Cux1
Tgif1
Nfat5
Foxq1
Kmt2a
Bptf
Prdm9
Arid3b
Usf2
Gtf2i
Rfx1
Rarg
Stat1
Ets2
Patz1
Stat6
Zfp398
Irf1
Foxo3
Maf
Stat2
Stat5b
Pou2f2
Nrf1
Rfx3
Adult
E2f1
Klf12
Esr1
E2f8
Foxm1
Mxd3
E2f2
E2f7
Hoxa7
Mycl
Ets1
Mbd2
Dnmt1
Pura
Foxd2
Ar
Ctcf
Arntl
Hoxa6
Clean
Mafb
Tfec
Nr1h3
Pax5
Hsf1
Zfp608
Spib
Ahr
Klf10
Zfp703
Snai1
Mafk
Pknox1
Nr1d2
Nr2c1
Zfp395
Etv5
TN, Adult
Stat5a
Yy1
Tcf4
Sox5
Klf1
TN, Clean
Klf7
Zfp143
Zbtb6
Zfp281
Tcf7
Irf3
Klf11
Thra
Srebf2
Adult, Clean
Clec4a1
Ebf1
Zfp366
Zfp385a
Pparg
Spi1
Pou2af1
Spic
Zfp422
TN, Adult, Clean
Sox4
Foxo1
Ikzf1
Egr3
Lef1
Sp3
Egr1
Egr2
Cell cycle and division
Short term effector and memory
Naive or late effector or memory
Short term effector or memory
Late effector or memory
Initial cytokine or effector response
Preparation for cell division
Naive and late memory
Early effector late memory
Memory precursor
Positive targets
$-\log_{10}$(adj p−val)
30
20
10
0
Negative targets
$-\log_{10}$(adj p−val)
20
10
0
Direction
Enriched on positive targets
Enriched on negative targets
$-\log_{10}$(p−val)
2.5
5.0
7.5
10.0
12.5

**Supplementary Fig. 4: Core TFs in different naïve CD8+ T cell subsets**

(A) GSEA of predicted targets of VM, neonatal and dirty core TFs on differential expression results between naïve CD8+ T cell subsets. (B) Hypergeometric test on predicted targets of VM, neonatal and dirty core TFs using gene sets from ImmGen delineate CD8+ T cells immune response. P-values smaller than $10^{-13}$ were plotted as $10^{-13}$. (C) GSEA of predicted targets of TN, adult and clean core TFs on differential expression results between naïve CD8+ T cell subsets. (D) Hypergeometric test on predicted targets of TN, adult and clean core TFs using gene sets from ImmGen delineate CD8+ T cells immune response. P-values smaller than $10^{-13}$ were plotted as $10^{-13}$. Related to Fig. 4.

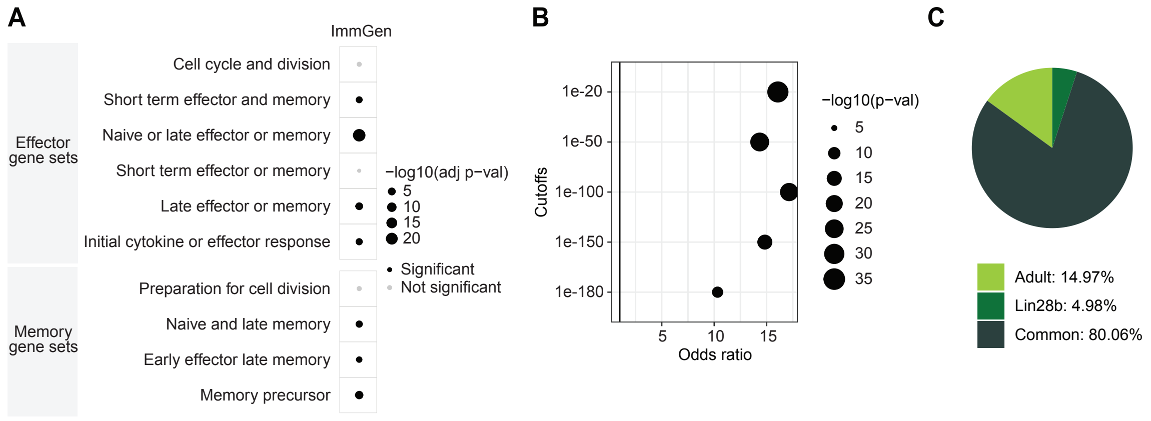


**Supplementary Fig. 5: *Eomes* binding at effector genes in neonatal cells**

(A) Enrichment analysis on genes associated with *Eomes* CUT&Tag peaks using ImmGen gene sets. (B) Enrichment of CUT&Tag-and-motif-analysis-derived *Eomes* targets (CUT&Tag peaks were restricted to those with at least one *Eomes* motif occurred) and network-derived *Eomes* targets at CUT&Tag peaks of different cutoffs. (C) Percentage of *Eomes* CUT&Tag Peaks with different binding in adult and Lin28b cells. Related to Fig. 5.

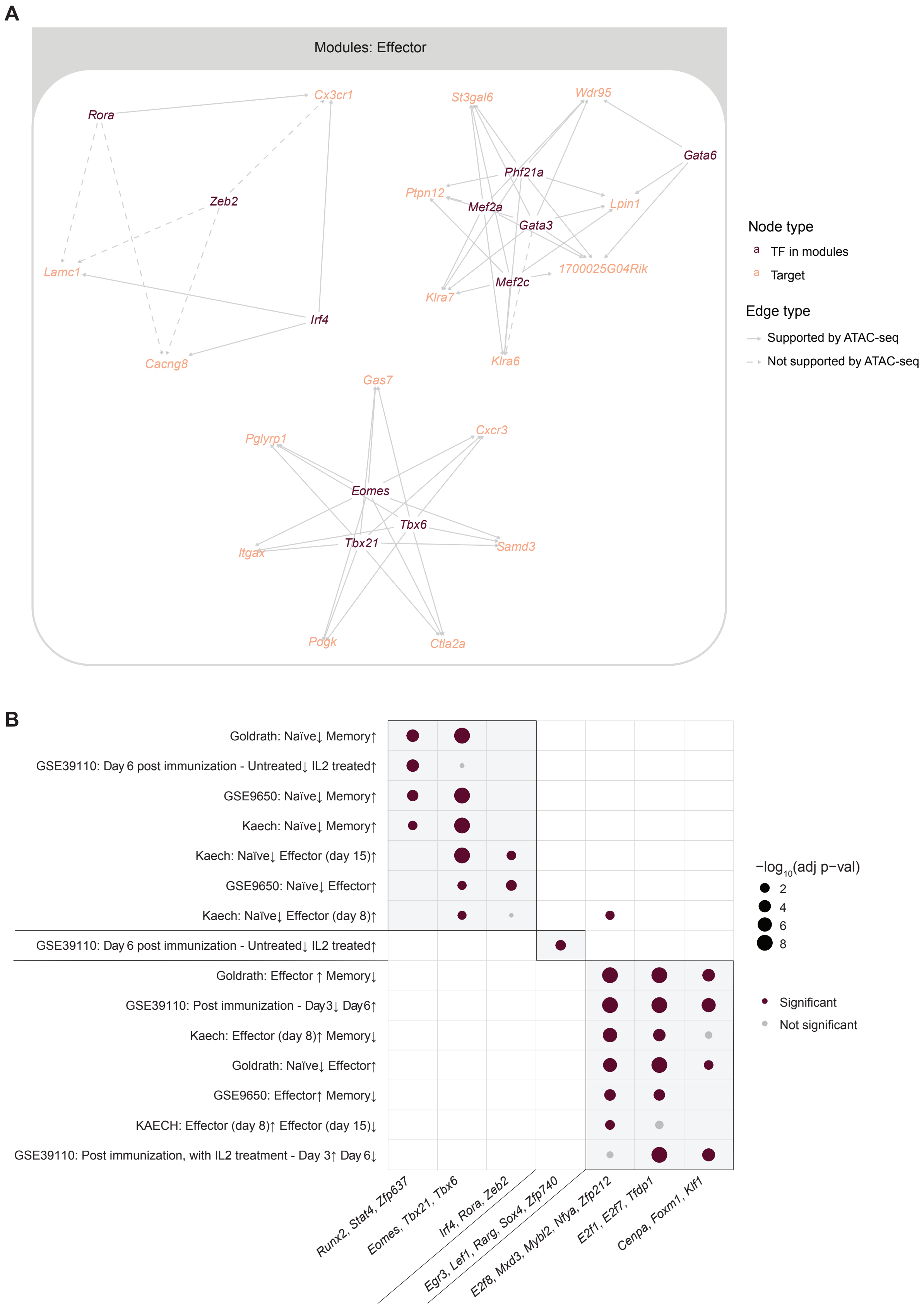
A
Modules: Effector
Rora
Cx3cr1
Zeb2
Lamc1
Irf4
Cacng8
St3gal6
Wdr95
Gata6
Phf21a
Ptpn12
Mef2a
Gata3
Lpin1
1700025G04Rik
Mef2c
Klra7
Klra6
Gas7
Cxcr3
Pglyrp1
Eomes
Tbx6
Tbx21
Itgax
Samd3
Pogk
Ctla2a
Node type
TF in modules
Target
Edge type
Supported by ATAC-seq
Not supported by ATAC-seq
B
Goldrath: Naïve↓ Memory↑
GSE39110: Day 6 post immunization - Untreated↓ IL2 treated↑
GSE9650: Naïve↓ Memory↑
Kaech: Naïve↓ Memory↑
Kaech: Naïve↓ Effector (day 15)↑
GSE9650: Naïve↓ Effector↑
Kaech: Naïve↓ Effector (day 8)↑
GSE39110: Day 6 post immunization - Untreated↓ IL2 treated↑
Goldrath: Effector ↑ Memory↓
GSE39110: Post immunization - Day3↓ Day6↑
Kaech: Effector (day 8)↑ Memory↓
Goldrath: Naïve↓ Effector↑
GSE9650: Effector↑ Memory↓
KAECH: Effector (day 8)↑ Effector (day 15)↓
GSE39110: Post immunization, with IL2 treatment - Day 3↑ Day 6↓
Runx2, Stat4, Zfp637
Eomes, Tbx21, Tbx6
Irf4, Rora, Zeb2
Egr3, Lef1, Rarg, Sox4, Zfp740
E2f8, Mxd3, Mybl2, Nfya, Zfp212
E2f1, E2f7, Tfdp1
Cenpa, Foxm1, Kif1
−log10(adj p−val)
2
4
6
8
Significant
Not significant

C

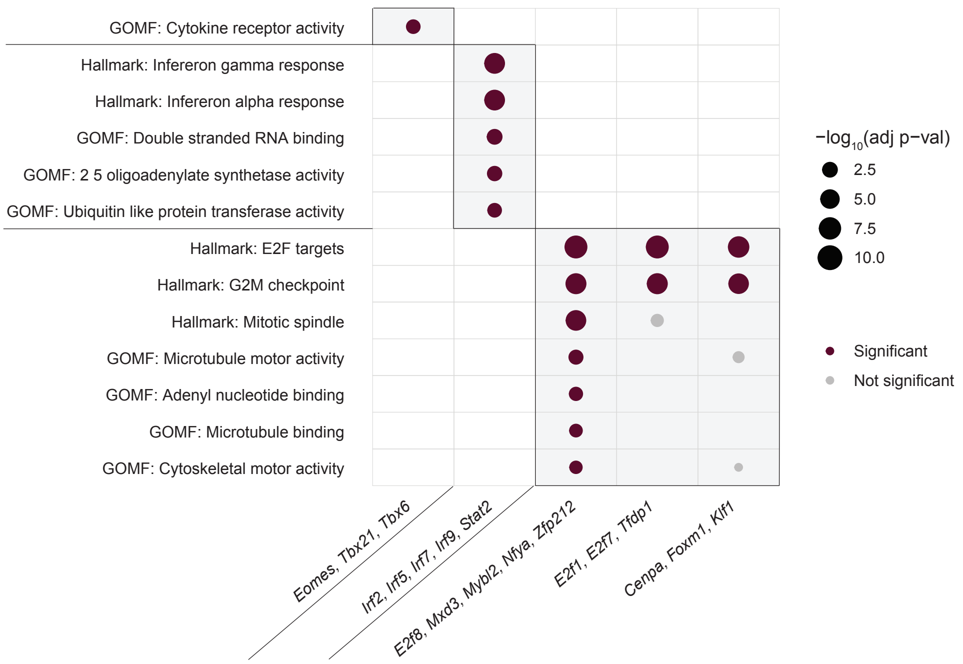
GOMF: Cytokine receptor activity
Hallmark: Infereron gamma response
Hallmark: Infereron alpha response
GOMF: Double stranded RNA binding
GOMF: 2 5 oligoadenylate synthetase activity
GOMF: Ubiquitin like protein transferase activity
Hallmark: E2F targets
Hallmark: G2M checkpoint
Hallmark: Mitotic spindle
GOMF: Microtubule motor activity
GOMF: Adenyl nucleotide binding
GOMF: Microtubule binding
GOMF: Cytoskeletal motor activity
Eomes, Tbx21, Tbx6
Irf2, Irf5, Irf7, Irf9, Stat2
E2f8, Mxd3, Mybl2, Nfya, Zfp212
E2f1, E2f7, Tfdp1
Cenpa, Foxm1, Klf1
−log10(adj p−val)
2.5
5.0
7.5
10.0
Significant
Not significant

D

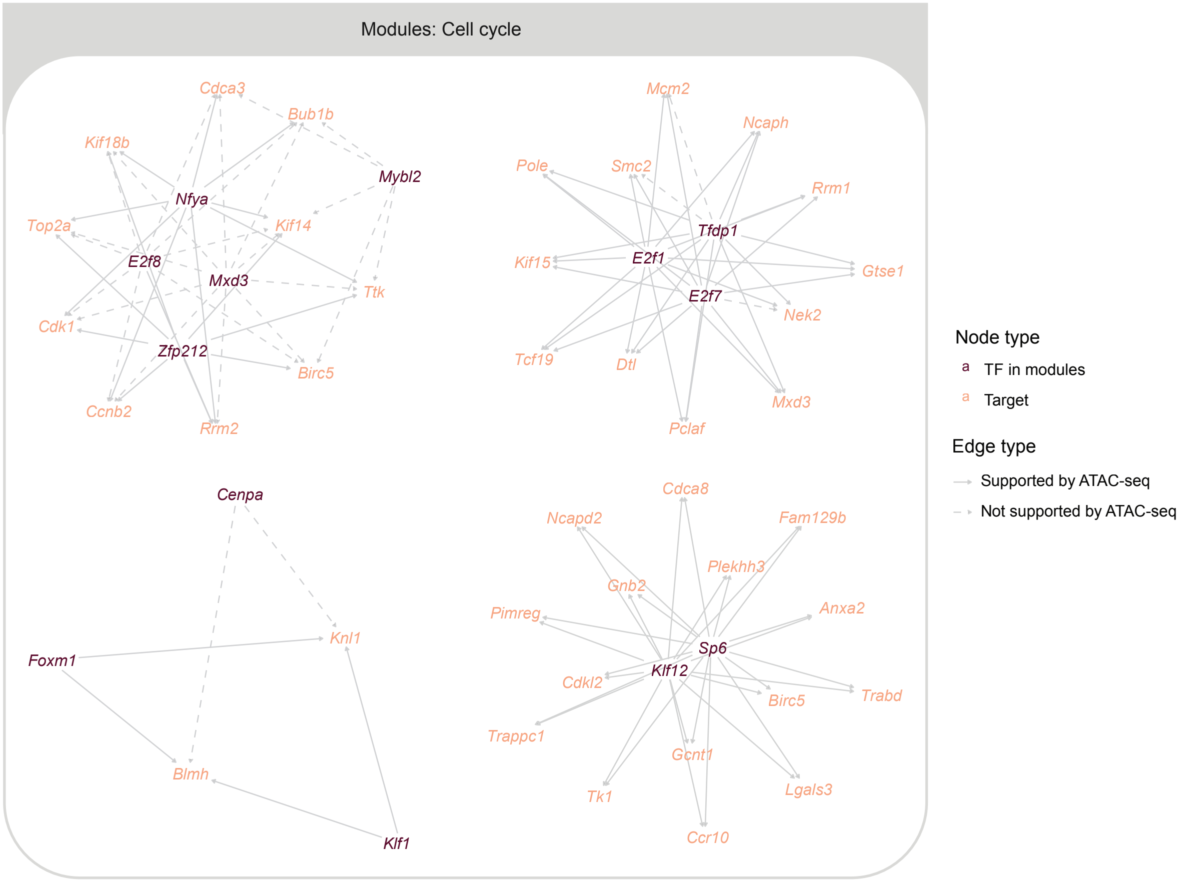
Modules: Cell cycle
Cdca3
Bub1b
Kif18b
Mybl2
Nfya
Top2a
Kif14
E2f8
Mxd3
Ttk
Cdk1
Zfp212
Birc5
Ccnb2
Rrm2
Mcm2
Ncaph
Pole
Smc2
Rrm1
Tfdp1
Kif15
E2f1
Gtse1
E2f7
Nek2
Tcf19
Dtl
Mxd3
Pclaf
Cenpa
Knl1
Foxm1
Blmh
Klf1
Cdca8
Ncapd2
Fam129b
Plekhh3
Gnb2
Anxa2
Pimreg
Sp6
Klf12
Cdkl2
Birc5
Trabd
Trappc1
Gcnt1
Tk1
Lgals3
Ccr10
Node type
TF in modules
Target
Edge type
Supported by ATAC-seq
Not supported by ATAC-seq

**Supplementary Fig. 6: Co-regulatory TF modules preprogram naïve cells for effector and memory functions**

(A) Subnetwork of modules with roles in effector functions (showing TFs and targets shared by more than 3, 4, 3 TFs respectively). (B) Hypergeometric test on predicted positive targets shared by >50% of TFs in each TF module using ImmuneSigDB gene sets. Adjusted p-values smaller than $10^{-8}$ were plotted as $10^{-8}$. (C) Hypergeometric test on predicted positive targets shared by >50% of TFs in each TF module using Hallmark and GO Molecular Functions gene sets. Adjusted p-values smaller than $10^{-8}$ were plotted as $10^{-8}$. (D) Subnetwork of modules with roles in regulating cell cycle (showing TFs and targets shared by more than 4, 3, 3, 2 TFs respectively). Related to Fig. 6.

**Supplementary Table 1**. TF expression levels across naïve CD8+ T cell subsets, scaled by rows (samples). Related to Fig. 1H.

**Supplementary Table 2**. Gene expression levels across naïve CD8+ T cell subsets, scaled by rows (samples). Related to Fig. 1I.
